# Melodic Modulation of Pain and Cognition: Neurobehavioral Effects of Indian Classical Music in Mice

**DOI:** 10.64898/2026.08.27.742492

**Authors:** Kanchan Mukherjee, Tuhin Bhattacharya, Sabnur Parvage, Sumoyee Ghosh, Rimi Das, Hemanta Mandal, Rakhi Dey Sharma, Sanjit Dey

**Affiliations:** Department of Physiology, University of Calcutta, 92 APC Road, Kolkata 70009, West Bengal, India; Department of Physiology, Rammohan College, 102/1, Raja Ram Mohan Sarani, Kolkata, West Bengal 700009; Centre for Nanoscience and Nanotechnology (CRNN), University of Calcutta, 92 APC Road, Kolkata 70009, West Bengal, India; Centre with Potential for Excellence in Particular Area (CPEPA), University of Calcutta, 92 APC Road, Kolkata 70009, West Bengal, India

**Keywords:** Pain, Non-Pharmacological Method, Indian Instrumental Music (IIM), Behavior, Neurotransmitters, Neuroplasticity, mRNA Expression

## Abstract

**Background:** Despite advances in pain management, effective analgesics in pain situations remain elusive. Opioids and non-opioids carry risks of neurotoxic and psychedelic effects with adverse physiological outcomes. Indian instrumental music (IIM) mitigates subacute pain by rewiring neurochemical synergy as an evidence-based, non-invasive, non-pharmacological system to mitigate pain.

**Objective:** Investigating therapeutic efficacy of IIM in mitigating subacute pain by analyzing behavioral, peripheral, and central neurochemical re-tuning.

**Methods:** Mice were divided into Control, Pain, Pain+Music, and Music groups. Pre-treatment behavioral parameters were compared with those observed after 14 days IIM exposure. Evaluations included nociceptive latencies (hot-plate/tail-flick), locomotion (Open Field Test), and anxiety (Elevated Plus Maze). Molecular analyses quantified peripheral neuropeptides (SP, NK-1R, CGRP), serum cortisol, spinal neurotrophic factor, neurotransmitters (glutamate, GABA, dopamine (DA), 5-HT), BDNF, and mRNA expression of BDNF, Ntrk1R/2R, and D1R in cortex, thalamus, hippocampus and hypothalamus. All procedures adhered to IAEC guidelines.

**Results:** IIM yielded 3.9-4.4-fold antinociceptive improvements, 3.3-fold locomotor restoration, and 3.6-4.9-fold anxiolysis. 14 days IIM exposure reduced peripheral nociceptive-neuropeptides 1.3-2.0-fold (SP, NK-1R, CGRP), serum cortisol 1.3-fold, and spinal glutamate, serotonin levels 1.5- and 1.3-fold. An enhanced expression of spinal GABA, DA about 1.5-fold, and BDNF by 1.3-fold was observed after music listening. Brain-region-specific differential mRNA-expression at cortex, thalamus, hypothalamus and hippocampus, revealed the neuromodulatory impact of rhythmic music in formalin-induced murine pain-model.

**Conclusion:** Gross reduction of pain parameters demonstrates therapeutic potential of IIM as multilevel neuromodulator to suppress the multidimensional stressor, pain, via peripheral desensitization, spinal E-I balance, and differential calibration of BDNF/Trk/D1R plasticity at specific brain-regions.

**GRAPHICAL ABSTRACT:** 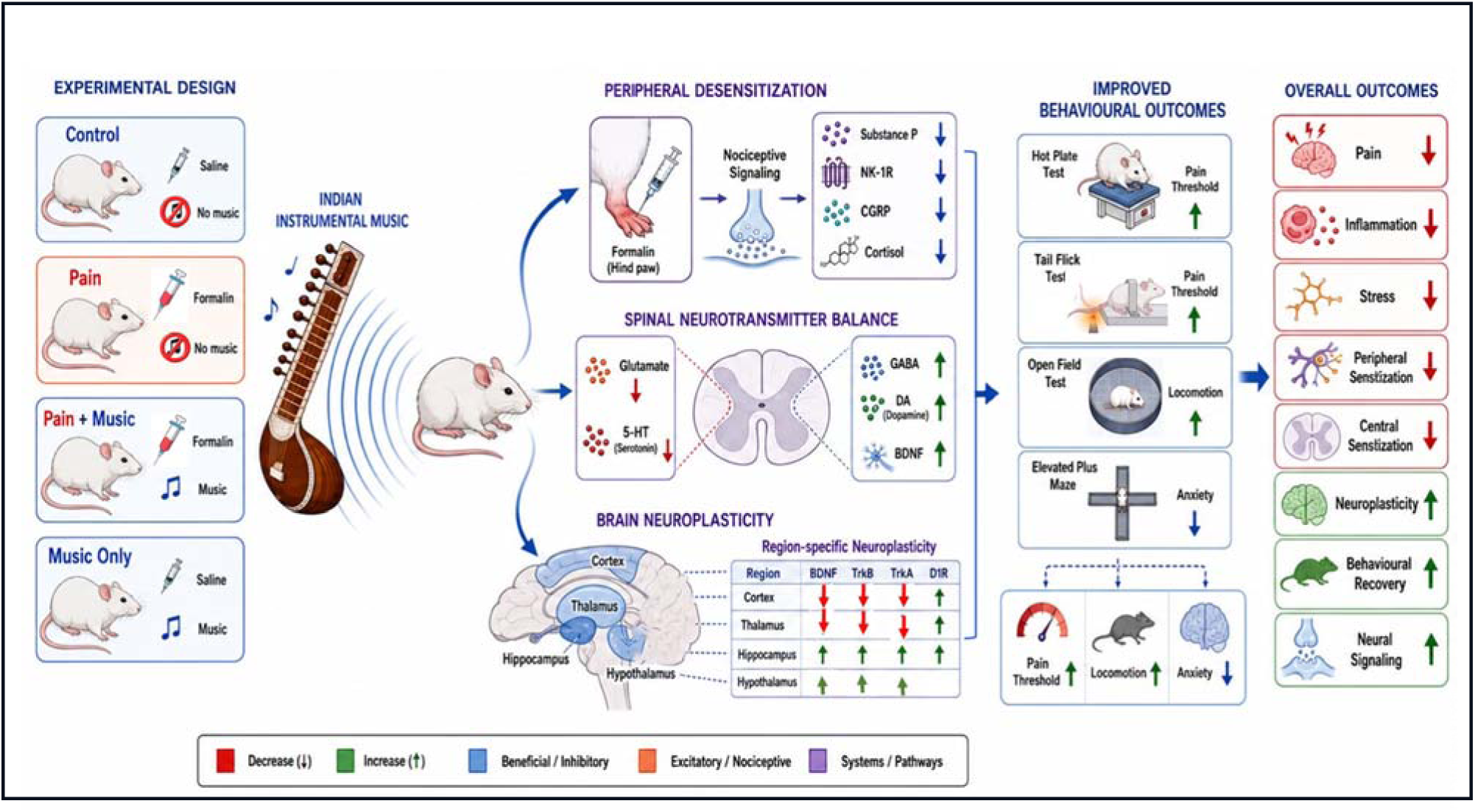

## 1. Introduction

Pain is a pervasive issue affecting millions worldwide, significantly impairing quality of life and posing substantial economic burdens. It is defined as an unpleasant sensory and emotional experience associated with, or resembling that associated with, actual or potential tissue damage burdens emerging from coordinated activity across peripheral, spinal, and supraspinal systems [1]. According to physiological function, pain is classified into nociceptive (physiological) and pathological pain. The pathological or clinical pain is divided into acute and chronic pain. Acute pain persists up to the usual time of recovery or healing [2]. Whereas in clinical practice, pain that lasts or recurs for longer than 3 months is defined as chronic pain [3]. And there is also a third category of pain, termed “subacute pain”; it is a longer-lasting subset of acute pain, persists between 6 weeks and 3 months [4].

From ancient times till now, it is our living instinct to replenish pain-stress with pleasure. Although conventional analgesics remain widely used, their limitations in long-term tolerability, side-effect, cost-burden, encouraged interest for search of non-pharmacological pain-management strategies. Music, the age-old universal emotion of mankind, becomes our saving grace which can holistically act across multiple levels of the pain network. Literature of music-based interventions further uphold the idea that music can influence emotional state and physiological arousal, engaging descending analgesic circuitry involving cortical, brainstem, and spinal regions and modifies pain-related experience. Recent studies are supporting music, as a promising adjunct for pain-management, with evidence pointing toward interactions between reward, stress, and nociception [5,6].

Despite music’s marked potency as neuropsychiatric modulator, proper mechanistic neurobiological evidence linking music exposure to transcriptional regulation within supraspinal pain networks are poorly understood. To, fulfill this gap we, developed a formalin-induced subacute-pain model and introduced 14 days Indian instrumental music (IIM) as treatment, to study whether pain-related behavior is shaped only by peripheral and spinal neurotransmission or also by region-specific gene expression in brain structures that integrate nociception, emotion, memory, and neuroendocrine responses. Hippocampus which contributes to stress adaptation and affective memory, and hypothalamus, which regulates autonomic and neuroendocrine responses, including hypothalamic-pituitary-adrenal axis activation are of esteemed relevance in this regard. The cortex and thalamus are also key nodes in sensory integration and descending modulation, making them biologically plausible targets for music-driven transcriptional reprogramming. [5,7–9]. We have investigated nociceptive behavior, locomotion, anxiety-like behavior, peripheral neuropeptides, systemic stress-markers, spinal neurotransmitters, and BDNF protein expression, along with supraspinal mRNA expression of BDNF, Ntrk1R, Ntrk2R, and D1R in cortex, thalamus, hippocampus, and hypothalamus. And observed that inflammatory pain was associated with elevated BDNF, Ntrk1R, and Ntrk2R expression in the pain group, whereas 14 days IIM exposure restored these changes, with region-specific normalization. Our findings support the hypothesis that rhythmic auditory stimuli act as a multilevel neuromodulator that may manage pain through peripheral desensitization, spinal excitatory-inhibitory rebalancing, and supraspinal transcriptional regulation. So, IIM may represent as a biologically active, non-invasive adjunct capable of modulating pain-related pathways from peripheral input to central gene expression.

## 2. Methods

### 2.1. Materials

The primary antibody and secondary IgG-HRP conjugated antibody were purchased from Cell Signaling Technology (Massachusetts, USA). All other reagents were of analytical grade and obtained from Sigma, St. Louis, MO or Merck SA (Darmstadt, Germany) or from local vendors.

### 2.2. Animals and Experimental Design

All experimental protocols and methods were carried out in accordance with relevant guidelines and regulations approved by the Institutional Animal Ethics Committee (IAEC), University of Calcutta, Kolkata, India;[IAEC-V/T/SD-14/KANCHAN MUKHERJEE/2024],which is registered Committee for Control and Supervision of Experiments on Animals (CCSEA), a statutory Committee of Department of Animal Husbandry and Dairying (DAHD), Ministry of Fisheries, Animal Husbandry and Dairying (MoFAH&D), Govt. of India we followed the standard Animal Research: Reporting of *in vivo* Experiments (ARRIVE) guidelines and Animal Research: Reporting of *in vivo* Experiments (ARRIVE). For this experiment, Swiss albino male mice (20–25 g) were procured randomly from a CCSEA registered supplier and maintained in standard environmental conditions (25±2 ° C, humidity 50±2%) maintaining 12 hr light/dark-cycle with food. Animals were acclimated for 1 week before testing. Animals were randomly assigned to four groups(n=3) for this study:

**Control:** Received normal saline (0.9% NaCl, 20μl) as vehicle control in the alternative hind paw on three specific treatment days, as it is the volume administered in the following groups [10].

**Pain:** Received 20μl, 2.5% formalin dissolved in normal saline v/v in the alternate hind paw for three specific treatment days.

**Pain+Music:** Analogous to the Pain group, but were exposed to 4 hours of daily IIM for 14 consecutive days from 18:00 to 22:00.

**Music:** Similar to the control group, additionally exposed to IIM like the Pain+Music group.

### 2.3. Music Selection

(Pain+Music) and Music group mice were exposed to 14 days of daily IIM for 4hours from 18:00 to 22:00. The music player was located 8 inches away from the mice cage, and IIM was played within a range of 47.4dBA to 74.6dBA. Regarding the selection, we designated the Indian classical instrumental tune “Raga Yaman”. The spectrum analysis revealed it as a combination of low frequency range (46-984Hz), to mid-frequency (10.312-20kHz) ranging up to high frequency (23.953kHz). This auditory stimulus falls within the murine audible range and overlaps with frequencies reported to be highly detectable in rodents, particularly in mid-to-high kilohertz band [11]. The primary musical information is centered in a pleasant mid-frequency range, with a gentle tapering of higher-frequency harmonics makes this music soothing rather than startling.

### 2.4. Open Field Test (OFT) and Elevated Plus Maze (EPM)

Mice were tested in a circular open field (OFT) and an elevated plus maze (EPM) at two different timepoints. Mice were habituated to the sound-proof experimental room (adjacent to the animal living room) for 1h before the start of behavioral tests. The environmental conditions including light-dark cycle, temperature, and humidity of the experimental room remained alike to the animal living room.

The behavioral patterns of mice were recorded in video using automated tracking software of ANY-Maze™ (Stoelting, USA). The track plots (in OFT) and heat maps (in EPM) generated in ANY-Maze™ software were used to clarify the pattern of mice exploration like ambulation (horizontal locomotor activities) in OFT and to clarify the pattern of mice exploration in EPM test. The occupancy plots generated in ANY-Maze™ software were used to signify the time stay in any zone of OFT and/or in the arms of EPM measuring the anxiety-like behavioral status. After every test, the experimental fields were carefully cleaned with 70% ethanol[10].

### 2.6. Determination of tail flick and paw withdrawal latency

To assess nociceptive responses, antinociceptive and antihyperalgesic activities of IIM tail flick and hot plate test was performed. To record the assess thermal hyperalgesia mice were placed on a tail flick analgesiometer with localized radiant heat to the tail at 52±1°C and tail flick latency was recorded as the time elapsed until the subject flicked its tail [12]. Hot plate test assessed the sensorimotor activity of rodents during painful stimulus by placing them on a metal test plate preheated at 52±1°C. Paw withdrawal latency was recorded as the time elapsed until the subjects licked or flicked its hind paw [13].

### 2.8. Immunoblot Assay

An equal amount of protein (40 μg) was loaded in each lane for 12% sodium dodecyl sulfate-polyacrylamide gel electrophoresis (SDS-PAGE) and transferred to a polyvinylidene difluoride (PVDF) membrane. The membrane was blocked with 5% skimmed milk (SM) solution overnight at 4°C. Immunoblotting was done using monoclonal antibodies to mouse BDNF protein. Beta-Actin was used as a loading control. Immunoblots were captured using a BioRadChemidoc (MP imaging system) then normalized and analyzed using Image J (NIH) software [10].

### 2.9. Neurotransmitter Assay

Glutamate, GABA, dopamine and serotonin levels from spinal L4–L6 segment-lysates were measured following the manufacturers protocols (MyBiosource, USA, catalog no. MBS2601720, MBS260709, MBS8807516, MBS1601042) and cortisol level was measured in serum (Calbiotech, USA, CO368S) [10].

### 2.10 Polymerase Chain Reaction

For mRNA expression studies, total RNA was isolated from brain regions using Trizol method (Sigma, St. Louis, MO). Then cDNA was prepared combining 1μg of RNA, using High-Capacity cDNA Reverse Transcription Kit (ThermoFisher 4368814) and RiboLock RNase Inhibitor (ThermoFisher EO0381), following manufacturer’s protocol. Applied Biosystems 2720, Thermal Cycler (Applied Biosystem, UK) was used for PCR amplification using resultant cDNA and gene-specific primers (Sigma Al). Ethidium bromide was used to stain the agarose gel after electrophoresis Image J.exe and Gel Doc EZ Imager (Bio-Rad) were used to evaluate the stained gels.

### 2.11. Data Management and Statistics

All data have been presented as mean±SEM. The significance of differences between the Pain+Music and Pain group was determined by the One-Way ANOVA following Tukey’s test using the software GraphPad Prism 8. A value of p< 0.05 was considered significant.

## 3. Results

### 3.1. Behavioral Study in Open Field Test (OFT)

#### 3.1.1. Restoration of Restricted Locomotor Activity after IIM Intervention

Locomotor activities, recorded in a circular open field, were suppressed in pain. Total distances travelled (DT, measured in m) was diminished after formalin treatment in the pain group (6.63±0.93), compared with the untreated control (23.60±1.21). DT was significantly restored in the Pain+music (21.63±1.40) group (Figure 2).

**Figure 1:**
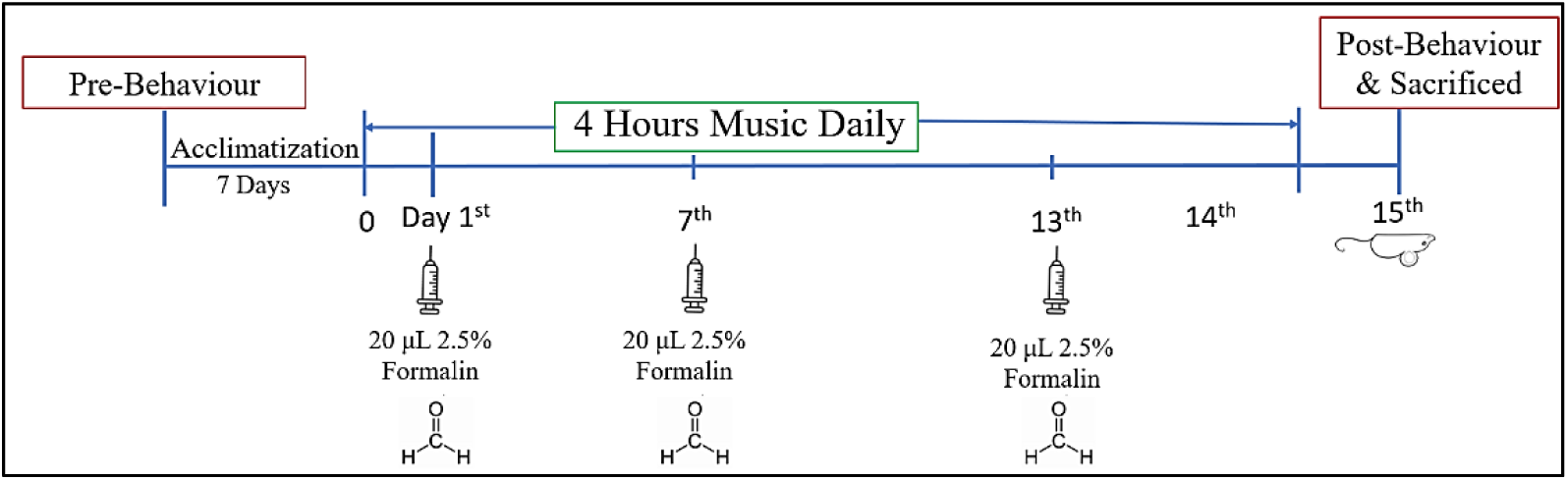
Study Design formalin-induced inflammatory-subacute pain model of mice exposed to 4 hours daily IIM for 14 days.

**Figure 2.1:**
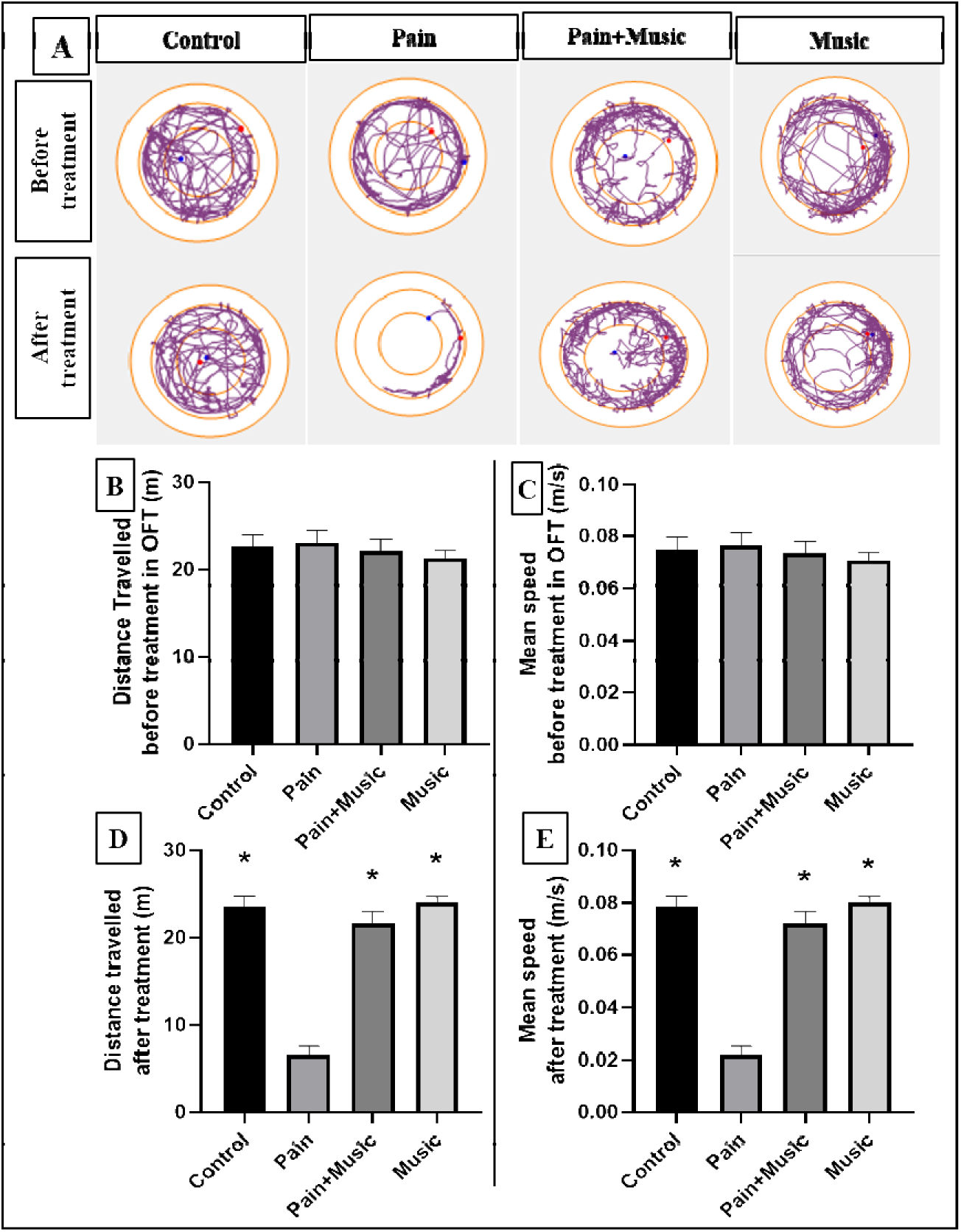
Locomotor activity in the open field. (A) Track plots showing locomotor activity before and after formalin and music treatment in a circular open field. Total distance travelled (DT, measured in meter) before (B) and after (D) formalin and music treatment. Mean speed (MS, measured in meter/second) before (C) and after (E) formalin and music treatment (n=3, p<0.05, *indicates significant difference with the formalin-treated group, Mean±SEM).

**Figure 2.2:**
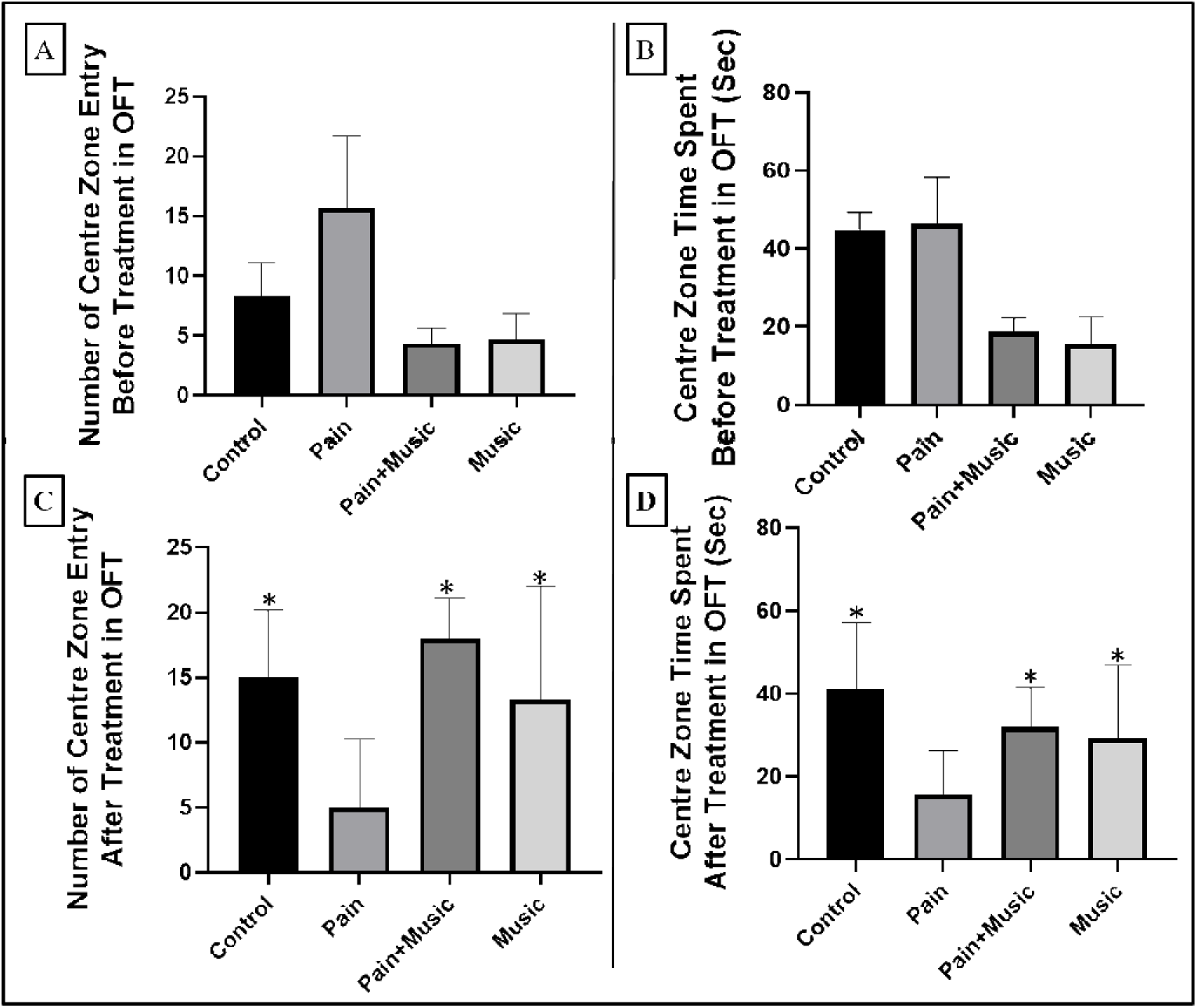
Anxiety and fear-like behavior in the open field: Total number of center zone entries before (A) and after (C) formalin and music treatment. Centre zone time spent (Seconds) before (B) and after (D) formalin and music treatment (n=3, p<0.05, *indicates significant difference with formalin-treated group, Mean±SEM).

**Figure 2.3:**
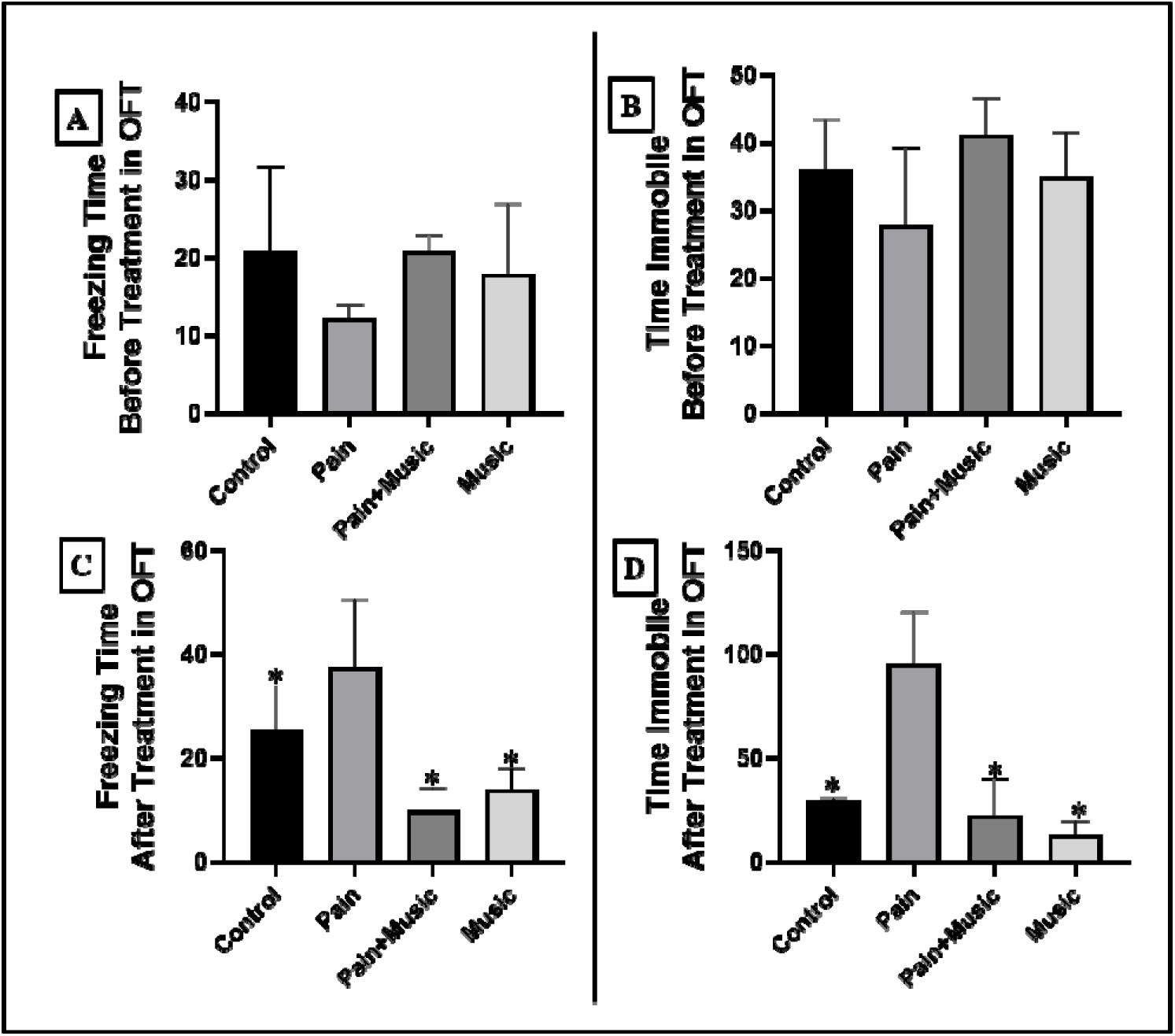
Anxiety and fear-like behavior in the open field. Total freezing time (Seconds) before (A) and after (C) formalin and music treatment. Total immobile time (Seconds) before (B) and after (D) formalin and music treatment (n=3, p<0.05, *indicates significant difference with formalin-treated group, Mean±SEM).

**Figure 3:**
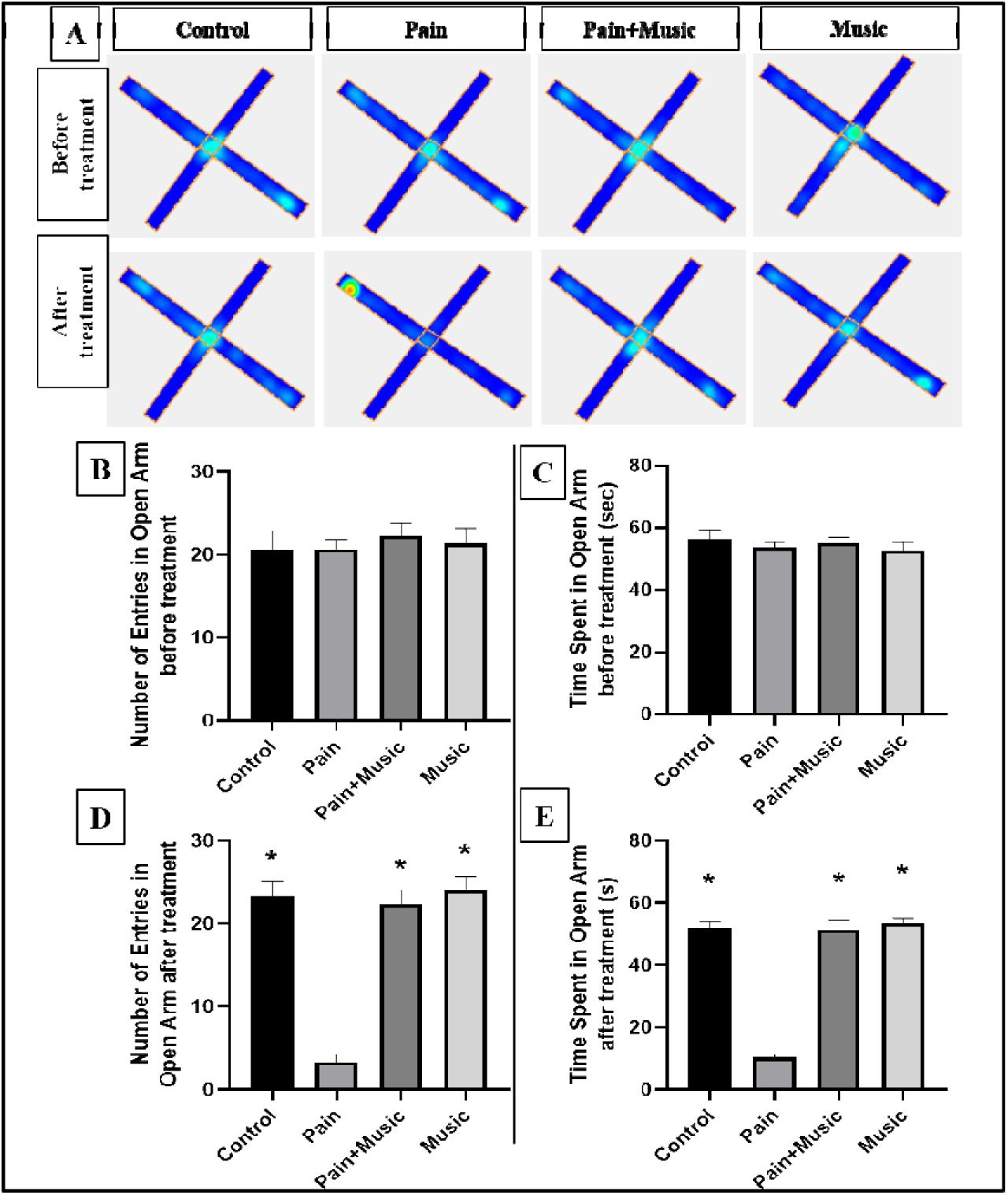
Anxiolytic phenomena of music in the elevated plus maze: (A) Heat map showing the presence of the rodents in the open arm and closed arm before and after formalin and music treatment in the elevated plus maze. Total number of entries in the open arm before (B) and after (D) formalin and music treatment. Time spent in the open arm (measured in seconds) before (C) and after (E) formalin and IIM treatment (n=3, p<0.05, *indicates significant difference with formalin-treated group, Mean±SEM).

**Figure 4:**
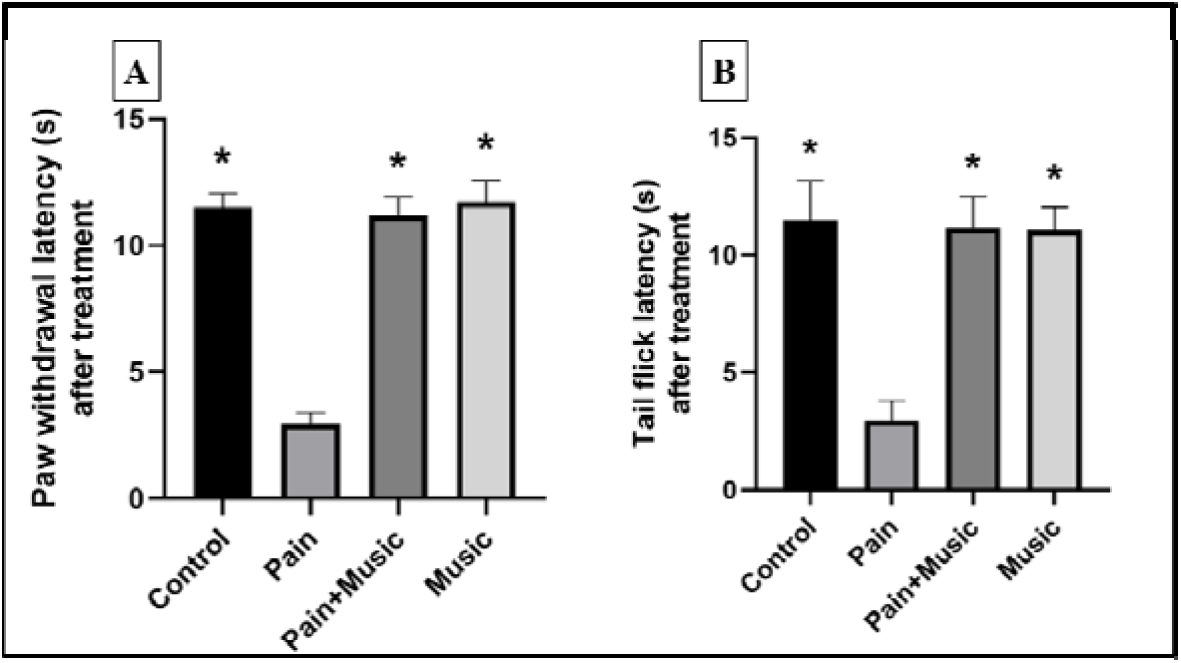
Pain sensitivity assessment (Hot plate test, Tail flick test). (A) Paw withdrawal latency (measured in seconds) after formalin and music treatment. (B) Tail flick latency (measured in seconds) after formalin and music treatment. (n=3, p<0.05, *indicates significant difference with formalin-treated group, Mean±SEM).

**Figure 5:**
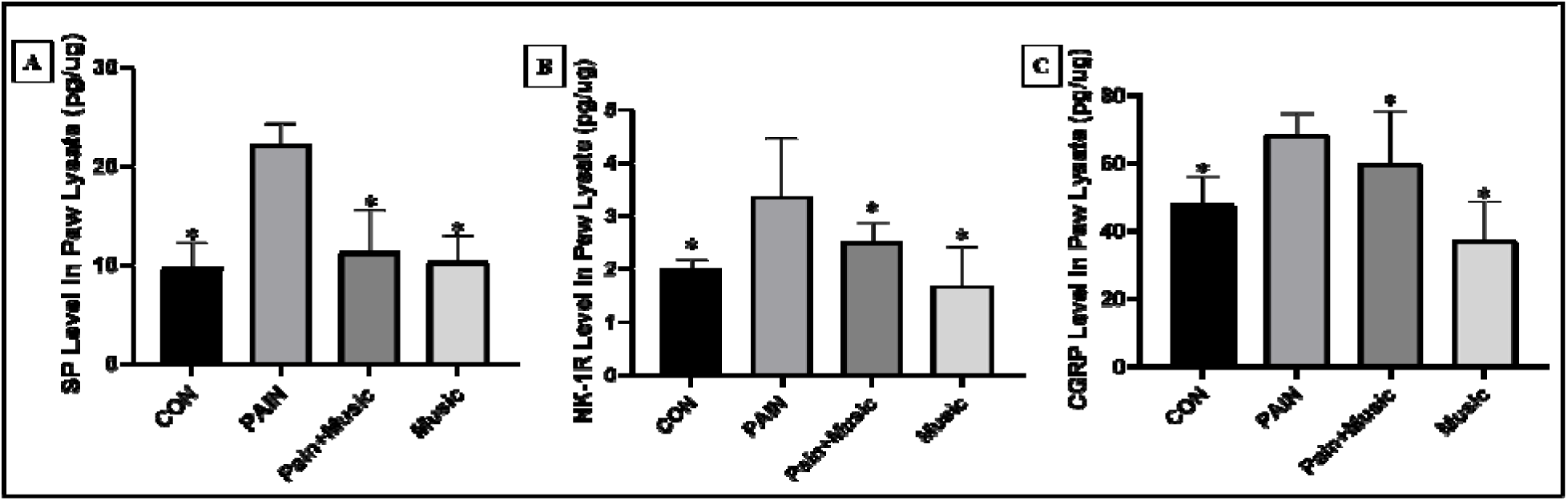
Excitatory neuropeptide in paw lysate. (A) Substance P (SP) (measured as pg/μg of protein) levels (B) Neurokinin-1 Receptor (NK-1R, measured as pg/μg of protein) levels (C) Calcitonin Gene Related Peptide (CGRP, measured as pg/μg of protein) levels after formalin treatment and music intervention (n=3, p<0.05, *indicates significant difference with the formalin-treated group, Mean±SEM).

Decreased mean speed (MS, meter/seconds), was observed in the pain group (0.022±0.003), compared to control (0.078±0.004) and restored in the Pain+music (0.072±0.004) group. Without any significant alteration in the music group (0.08±0.002) (Figure 2). The track-plots analysis further confirms IIM-intervened restoration of restricted movements.

#### 3.1.2. Recovery from Anxiety and Fear-like Behavior as found in OFT

To evaluate the modulation of behavioral and physiological signs of fear and anxiety-like behavior, total number of entries and time spent (in seconds) in the center zone, total immobile and freezing time (in seconds) in OFT were measured before and after treatments [14].

A significant decrease in the total number of center zone entries was recorded in pain group (5.00±1.75) compared to control (15.00±1.73), and an increase in the same was observed after IIM listening in the Pain+music group (18.00±1.04). The total time spent in the center zone was significantly reduced, after formalin treatment (15.70±2.05) compared to the untreated control group (41.10±3.10), and restoration of the same was recorded in the Pain+music group (18.00±1.04).

No significant alteration in total number of center zone entries or spent time at center zone of the circular open field was shown in the music group (13.33±2.89; 29.23±3.40) compared to control. Significant increase in total freezing time was recorded in pain group animals (37.57±3.07 sec) compared to the controls (25.57±1.98 sec), and was significantly calibrated after IIM exposure in the Pain+music group (9.87±1.04 sec).

Raise in time immobile was observed in the pain group (95.47±5.07 sec) compared to the control rodents (29.27±0.04 sec), and was reinstated significantly after IIM listening in Pain+music group (22.50±4.15 sec).

### 3.2. Elevated Plus Maze Test (EPM) depicts anxiolytic phenomena by IIM

To assess anxiety-responses in elevated plus maze test (EPM) number of entries (NE) and time spent (TS, second) in open arm (OA) was monitored before and after treatment[15]. Heat maps showed a confined position of animal in closed arm after formalin administration, whereas music induction retained the movement of animals. Significant lowering in number of OA entries was recorded after formalin treatment (3.33±0.88), with compare to the Control (23.33±17.64). NE was found to be significantly restored in the Pain+music (22.33±17.64) group.

Total time spent (TS, in seconds) in OA was significantly lowered in pain group (10.43±1.01), with compare to control (51.90±1.90). Significant restoration and upregulation of TS, in the Pain+music (51.50±3.05) animals was recorded.

No significant alteration was found in the music group’s NE (24.0±17.32) or TS (53.50±1.68) with respect to the control. Hence, rhythmic melody introspection demonstrates anxiolysis and exploratory activity in formalin-induced murine pain model.

### 3.3. Antihyperalgesic and Antinociceptive Effects of Rhythmic Music

To determine IIM’s antinociceptive efficacy on heat hyperalgesia, tail flick latency (TFL) and paw withdrawal latency (PWL) both in seconds, were measured in Tail-flick and Hot-plate analgesiometer. TFL manifests sensitivity to noxious stimuli, where flicking tail away from the heat source (16) denotes nociceptive response elicited by spinal flexion reflex. Formalin significantly lowered TFL (2.96±0.84), compared to the control group (11.50±1.67) as a prominent index of heat hyperalgesia. Whereas post IIM treatment TFL upregulated(11.17±1.34) significantly as an anti-hyperalgesic effect. Formalin treatment heightened thermal hyperalgesia, reducing PWL (2.96±0.41) significantly with comparison to the untreated control (11.50±0.56); and upregulated significantly after music intervention in the Pain+music group (11.17±0.75).

TFL(11.07±0.98) and PWL(11.73±0.84) of Music (11.73±0.84) group demonstrated no significant alterations. Unlike TFL, PWL results are dependent on the rodents’ motor coordination, so its upregulation indicates better motor coordination[16].

### 3.4 Peripheral Modulation of Inflammatory Nociception in Paw Tissue

In peripheral paw tissue-homogenate nociceptive undecapeptide Substance P (SP) (pg/µg) was significantly elevated in the pain group (22.16 ± 0.5433) compared with control mice (9.718 ± 0.6353), but it markedly reduced to near-control values, in the Pain + Music group (11.29 ± 1.064).

Consistent with enhanced SP signaling, expression of its high-affinity receptor, neurokinin-1 receptor (NK-1R), was significantly increased in pain (3.367 ± 0.3190 pg/µg) relative to controls (2.005 ± 0.04725 pg/µg). Music intervention significantly reversed NK-1R expression in the Pain+Music group (2.511 ± 0.1009 pg/µg).

Calcitonin gene-related peptide (CGRP) was significantly elevated in periphery following formalin-treatment (68.60 ± 1.765) compared with controls (47.93 ± 2.365). In Pain + Music group its expression(50.09 ± 1.349), was toward control animals. Notably, music-only group exhibited a reduction in SP(10.35 ± 0.6599); NK-1R(1.687 ± 0.2113 pg/µg), and CGRP(37.45 ± 3.273), indicating IIM exert a suppressive effect on basal peripheral nociception.

### 3.5. Alteration in Spinal Neurotransmitter Level

To determine whether IIM can exert any modulatory effects on spinal excitatory-inhibitory neurotransmitters, their levels were evaluated from the L4–L6 spinal lysate(10). Significantly higher glutamate level was observed in formalin-treated group (2.29±0.10 vs. control 0.96±0.07). And was lowered in the pain+music (1.64±0.05) group, with no significant alteration in the music group (1.02±0.49) (Figure 7). Inhibitory neurotransmitter GABA was diminished in pain group (3.92±0.14 vs. control 6.29±0.10). GABA-level significantly upregulated in pain+music (5.73±0.04) group, the music group (6.36±0.08) did not show such alteration. The key neurotransmitter of reward pathway dopamine, was found to be diminished in pain(1.50±0.07 vs. control 2.66±0.09). Introducing IIM significantly recalibrated this spinal dopamine-level, in the pain+music (2.20±0.10) group, music group (2.63±0.16) remain devoid of any such alterations (Figure 6). Significant elevation in spinal-serotonin level was found in formalin-treated group (2.26±0.09 vs. control 1.43±0.05). It was found to be significantly depressed in pain+music (1.76±0.05) group, without any variations in the music-only group (1.39±0.12) (Figure 7). IIM induced overall exaggeration of inhibitory and reward-processing neurotransmitter at spinal-level.

**Figure 6:**
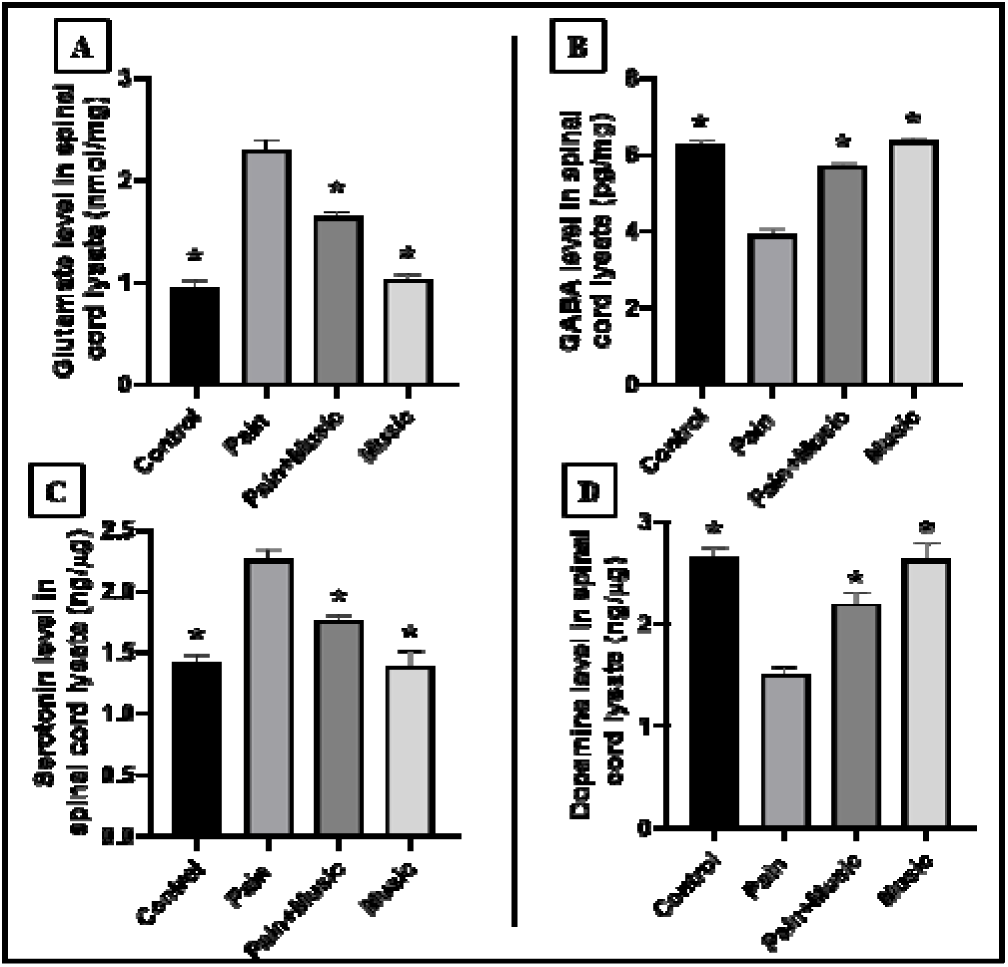
Excitatory and inhibitory neurotransmitters in spinal L4–L6 segment lysate. (A) Glutamate levels (measured as nmol/mg of protein) after treatment. (B) γ-Aminobutyric acid (GABA, measured as pg/mg of protein) levels after treatment. (C) Serotonin (measured as ng/μg of protein) levels after formalin and music treatment. (D) Dopamine levels (measured as ng/μg of protein) after treatment. (n=3, p<0.05, *indicates significant difference with the formalin-treated group, Mean±SEM).

**Figure 7:**
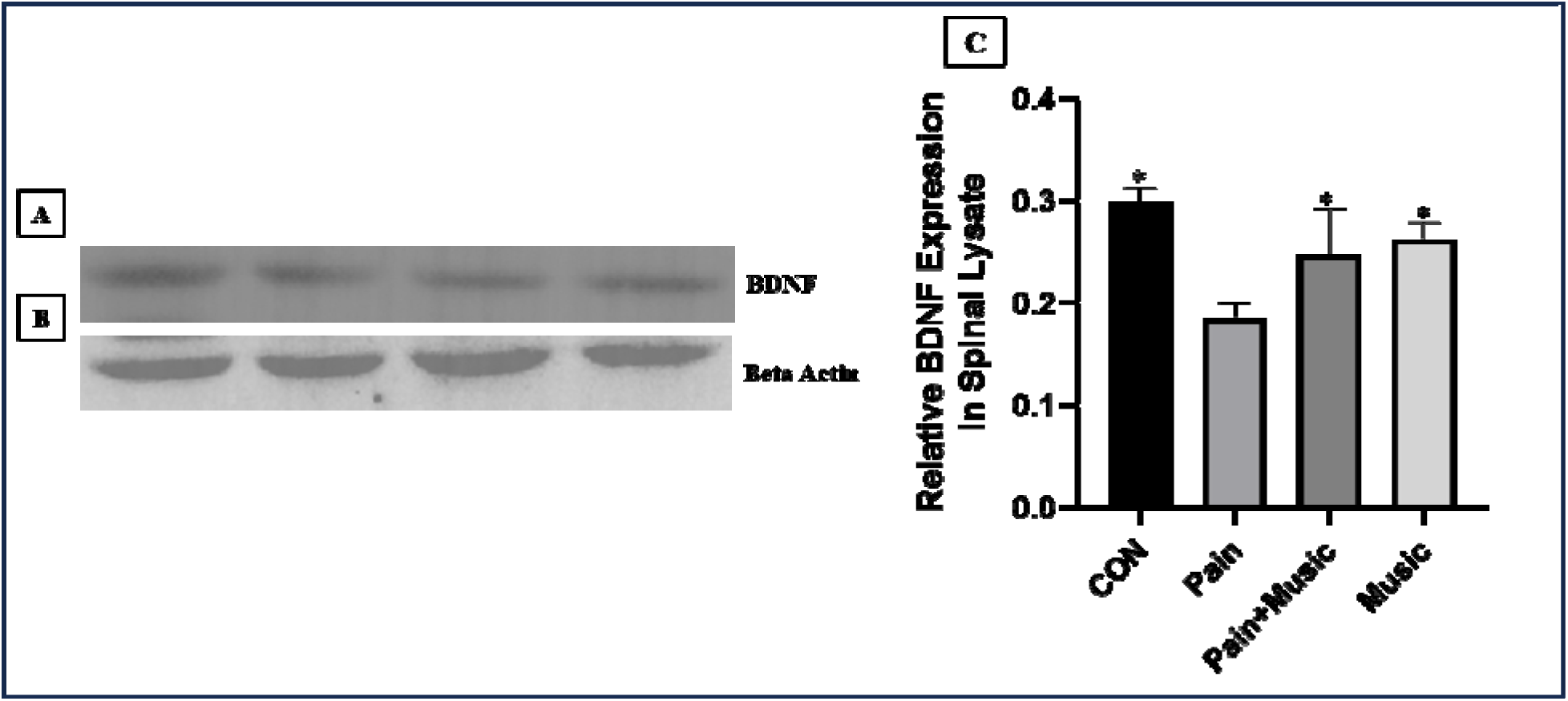
Cropped immunoblot image of BDNF (A) and Beta Actin (B); (C) Evaluation and densitometric analysis of Brain-derived Neurotrophic Factor (BDNF) in the spinal L4–L6 region after formalin and music treatment via immunoblot (n=3, p<0.05, *indicates significant difference with the formalin-treated group, Mean±SEM).

**Figure 8.1:**
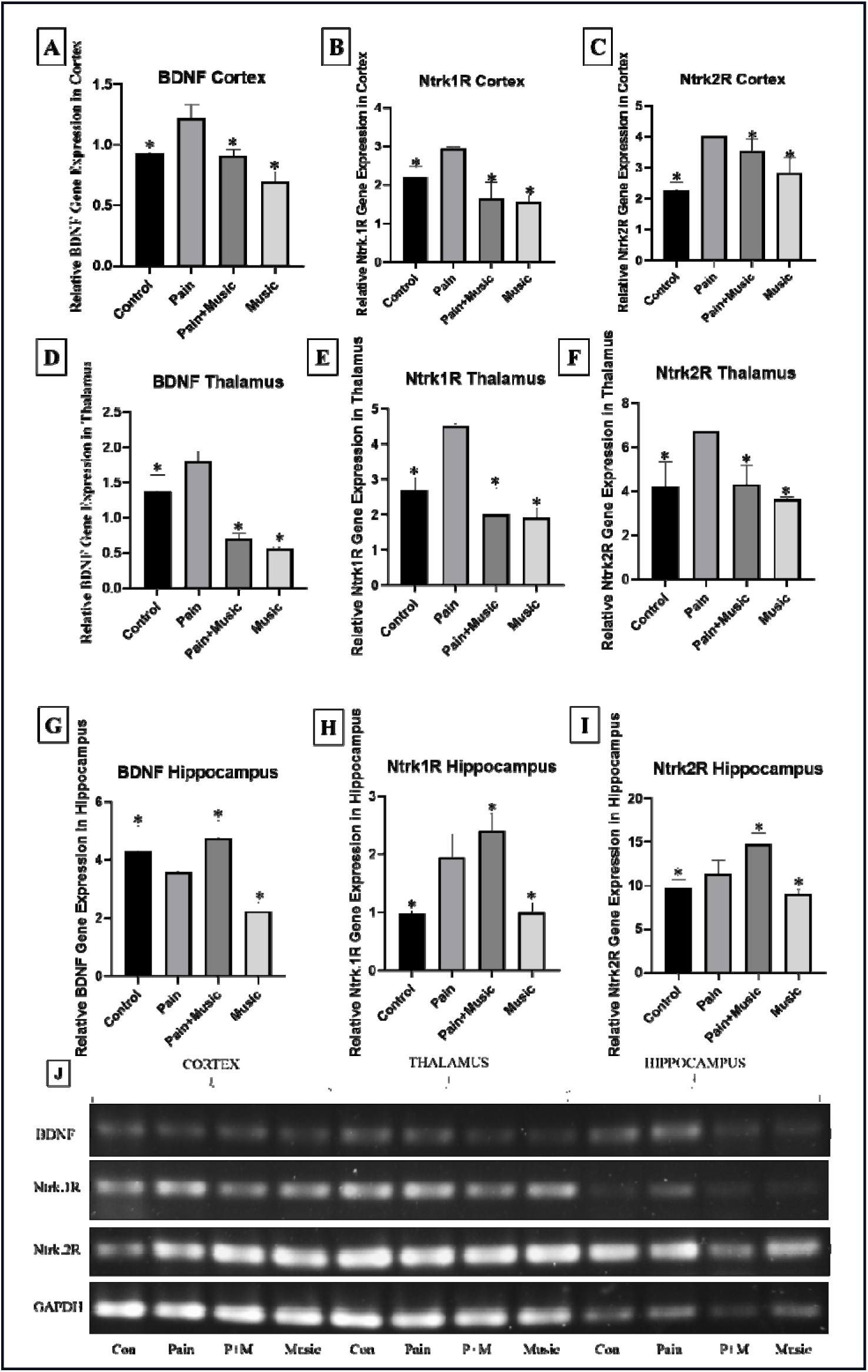
GAPDH normalized mRNA expression of BDNF (A), Ntrk1R(B), Ntrk2R (C) in the Cortex. GAPDH normalized mRNA expression of BDNF (D), Ntrk1R(E), Ntrk2R (F) in the Thalamus. GAPDH normalized mRNA expression of BDNF (G), Ntrk1R(H), Ntrk2R (I) in the hippocampus. (n=3, p<0.05, *indicates significant difference with the formalin-treated group, Mean±SEM). (H) Cropped agarose gel image.

**Figure 8.2:**
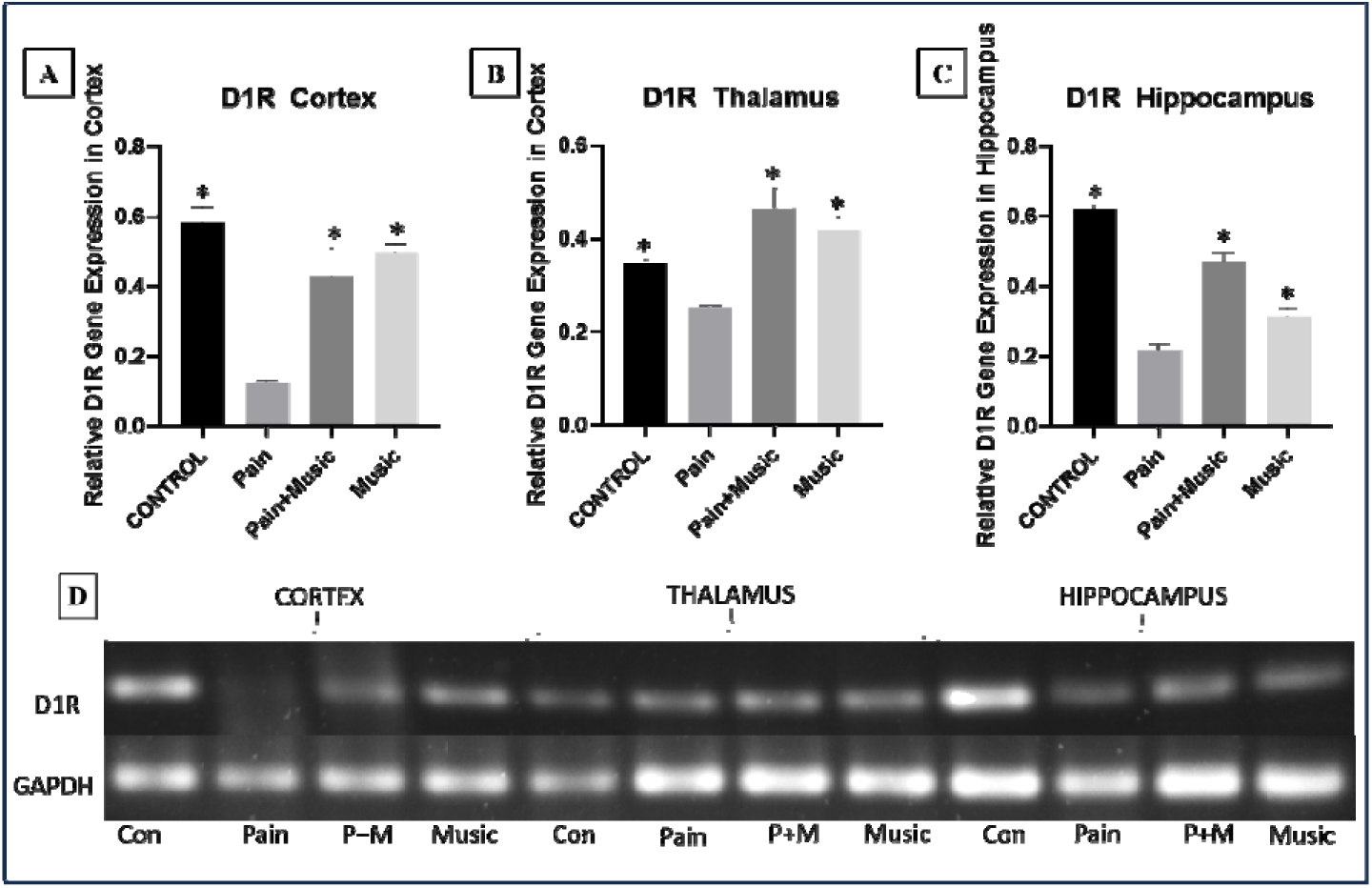
GAPDH normalized mRNA expression of Dopamine 1 receptor (D1R) Cortex (A), Hippocampus(B), Thalamus(C). (n=3, p<0.05, *indicates significant difference with formalin-treated group, Mean±SEM). (D): Cropped agarose gel image.

**Figure 9:**
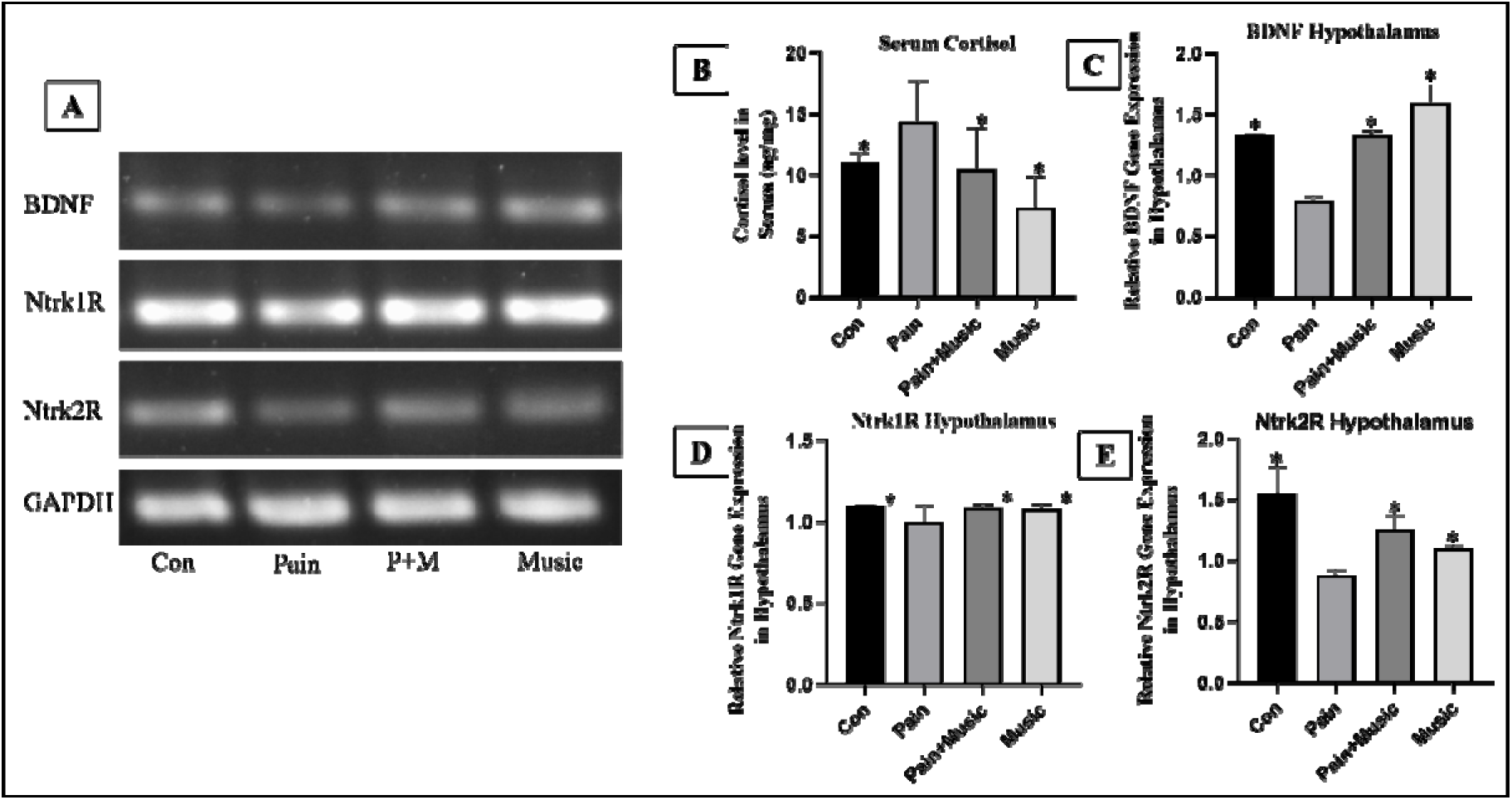
A. Cropped agarose gel image. B. Alteration in serum cortisol level across the groups. GAPDH normalized mRNA expression of BDNF (C), Ntrk1R(D), Ntrk2R (E) in the Hypothalamus. (n=3, p<0.05, *indicates significant difference with formalin-treated group, Mean±SEM).

### 3.6. Alteration in the Spinal Level of Brain-Derived Neurotrophic Factor (BDNF)

Spinal (L4-L6) immunoblot analysis showed, formalin administration reduced BDNF protein-expression (0.18±0.004 vs. control 0.29±0.004). However, pain+music (0.24±0.014) group has significantly restored it. Music group (0.26±0.005) also showed a significant increase in the spinal BDNF level (Figure:8).

#### 3.7.1 Supraspinal mRNA Expression of BDNF, Ntrk1R, Ntrk2R

The mRNA expression levels of brain-derived neurotrophic factor (BDNF), neurotrophic tyrosine kinase receptor 1 (Ntrk1R/TrkA), neurotrophic tyrosine kinase receptor 2 (Ntrk2R/TrkB) and Dopamine 1 receptor (D1R) were quantified via conventional PCR across the groups, and normalized with GAPDH.

##### 3.7.1.A Cortical Expression

BDNF mRNA expression increased significantly in the pain group (1.219 ± 0.039 vs. controls 0.932 ± 0.003), with reversal in the pain + music group (0.912 ± 0.016) and further reduction in the music-only group (0.695 ± 0.026). Ntrk1R expression rose in pain (2.946 ± 0.012 vs. controls 2.205 ± 0.091), decreasing in both pain + music (1.645 ± 0.143) and music-only (1.558 ± 0.055) groups. Ntrk2R followed suit, elevating in pain (4.014 ± 0.012) from controls (2.271 ± 0.087), then declining in pain + music (3.535 ± 0.127) and music-only (2.823 ± 0.168) conditions (Figure:9.1: A, B, C).

##### 3.7.1.B Thalamic Expression

Thalamic BDNF mRNA rose significantly in pain (1.801 ± 0.045 vs. controls (1.370 ± 0.080), reversing in pain + music (0.699 ± 0.026) and dropping further in music-only (0.557 ± 0.012). Ntrk1R increased in pain (4.513 ± 0.029 vs. controls 2.691 ± 0.115), with reductions in pain+music (2.003 ± 0.248) and music-only (1.922 ± 0.085) groups. Ntrk2R expression paralleled this, rising in pain (6.740 ± 0.145 vs. controls 4.230 ± 0.369), then falling in pain + music (4.316 ± 0.293) and music-only (3.630±0.034) groups (Figure:9.1: D,E,F).

##### 3.7.1.C Hippocampal Expression

Hippocampal BDNF mRNA decreased markedly in pain (3.587 ± 0.028 vs. controls 4.296 ± 0.296), with robust reversal in pain + music (4.749 ± 0.212) and further reduction in music-only (2.233 ± 0.109). Ntrk1R increased in pain (1.947 ± 0.133 vs. Controls 0.973 ± 0.018) and rose further in pain + music (2.412 ± 0.102), while music-only (0.995 ± 0.059) showed no significant change versus controls. Ntrk2R elevated in pain (11.36±0.516) versus controls (9.695±0.332), increased further in pain + music (14.77±0.421), and remained unchanged in music-only (9.030 ± 0.180) relative to controls (Figure:9.1: G,H,I).

#### 3.7.2 Supraspinal mRNA Expression of D1R

##### 3.7.2.A Cortical Expression

Cortical D1R mRNA expression observed to be decreased significantly in the pain group mice (0.126 ± 0.001 vs. control group (0.582 ± 0.026), with reversal in the pain + music group (0.426±0.048) and further increase in the music-only group (0.496 ± 0.026) (Figure:9.2: A).

##### 3.7.2.B Thalamic Expression

D1R mRNA expression in the thalamus decreased markedly in the pain group mice (0.253±0.001 vs. control group (0.346±0.006) and a robust reversal was observed after music intervention (0.464±0.025). Thalamic D1R mRNA expression level in the only music group (0.418±0.016), observed higher than the control group (Figure:9.2: B).

##### 3.7.2.C Hippocampal Expression

D1R level in the hippocampus was observed to be lowered in the pain condition (0.215±0.012 vs. control group mice (0.622± 0.004) and was significantly increased after music intervention (0.468 ± 0.015). The D1R mRNA expression level in only music group (0.314±0.012) was higher than the pain group, in hippocampus (Figure:9.2: C).

### 3.9 Normalization of HPA hyperactivity and restoration of hypothalamic BDNF/Trk expression in subacute inflammatory pain

Formalin-induced pain activated HPA-axis and concurrent reductions in hypothalamic neurotrophic signaling. Serum cortisol rose in the Pain group (14.38 ± 1.10 ng/mg) versus controls (10.99 ± 0.26 ng/mg), indicating pain-driven stress hormone dysregulation; exposure to IIM during the subacute phase restored cortisol to near-control levels in pain+music animals (10.48 ± 1.11 ng/mg) and provided a less-stress condition for music-only animals (7.32 ± 0.83 ng/mg) (Fig. 10). Parallel changes were observed when hypothalamic BDNF mRNA decreased markedly in pain (0.79 ± 0.025 vs. controls 1.33 ± 0.004), with robust reversal in pain + music (1.33 ± 0.024) and further increase in music(1.59 ± 0.085) groups. Ntrk1R decreased slightly with pain (0.99 ± 0.03 vs. Control 1.09 ± 0.001) and recovered in Pain + Music (1.08 ± 0.006), while Ntrk2R was markedly reduced by pain (0.88 ± 0.01 vs. Control 1.55 ± 0.07) and partially rescued by IIM (1.25 ± 0.04); Music-only levels approximated controls for both receptors (Ntrk1R 1.08 ± 0.008, Ntrk2R 1.10 ± 0.008).

## Discussion

Pain has long been tried to manage through approaches beyond pharmacology, especially to find an effective, affordable, long-term-usable, systemic adverse effects-free holistic analgesic strategy. Among non-pharmacological approaches, music has emerged as a promising candidate among neuromodulatory interventions, as it can influence affective state, stress responsiveness, and nociception, yet its underlying molecular and neurobiological basis remains incompletely defined[17].

To address this gap, we developed a formalin-induced subacute murine-pain model and examined the effects of 14 days daily IIM exposure. Since formalin evokes pain through peripheral inflammatory mediators followed by spinal [18] and supraspinal responses, this model offered an useful framework to investigate whether music can modulate pain across multiple levels of the neuroaxis. Although auditory stimulation in rodents may initially appear as subjective sensory input[11], our evidence suggests that rhythm-based sound can engage biologically meaningful behavioral and neural responses, making music a plausible contender for pain management. 14 days of IIM exposure produced a broad antinociceptive, anxiolytic, and locomotor-restoring effect. Importantly, these behavioral improvements were accompanied by modulation in coordinated cascade spanning peripheral nociceptive signaling, spinal-neurotransmitter balance, and supraspinal gene regulation. Pain is not merely a sensory experience but a multidimensional stressor that challenges both physiological and psychological homeostasis. Subacute nociceptive input activates peripheral inflammatory pathways, while engaging the hypothalamic–pituitary–adrenal (HPA) axis, elevating cortisol and sympathetic activity. At the central level, pain induces maladaptive neuroplastic changes across cortical, thalamic, hippocampal, and hypothalamic circuits, mirroring the effects of stress exposure. Thus, pain can be conceptualized as a dual stressor — simultaneously provoking inflammatory and neuroendocrine responses while burdening emotional and cognitive systems. Our findings suggest, IIM acted across this entire axis, reduced emotional distress and simultaneously normalizes sensory nociception and pain-related plasticity.

At the behavioral level, the pain group showed reduced locomotion, diminished exploratory drive, increased immobility/freezing, and marked impairment of thermal nociceptive thresholds, indicating that formalin suppressed both motor engagement and exploratory drive while enhancing defensive behavior. These changes are consistent with the view that inflammatory pain affects not only peripheral nociception but also recruits distributed central-neural systems that encode threat, aversion, and stress. The restoration of mean speed and total distance traveled in OFT (Figure:2.1) after IIM-exposure, indicates that music has intervened, not only pain burden but also the motivational and affective inhibition that accompanies sustained inflammatory nociception. Rhythmic auditory stimulation is known to engage reward-related and motor circuits, including the mesolimbic dopaminergic system, basal ganglia, and sensorimotor pathways, which together can support movement initiation and behavioral vigor [19,20].Thus, the locomotor recovery observed here likely reflects reactivation of circuits that integrate motivation, movement, and affective state. IIM improved anxiety-like phenotypes by restoring central zone activity, lowering immobility and freezing (Figure:2.2;2.3), increasing number of entries and time spent in open arms [21–24] indicates a shift away from defensive avoidance and toward exploration (Figure:3). These behaviors especially designate that animals are not merely experiencing nociception but perceiving environmental threat. Insula, hippocampus, amygdala, prefrontal cortex, and hypothalamus are key structures in this response, integrating nociceptive input with emotional appraisal and stress output [7,8,21,25].

This behavioral recovery is also reflected in the thermal nociception assays. Formalin pain reduced both paw withdrawal latency (PWL) and tail flick latency (TFL), confirming enhanced heat hyperalgesia and a lowered pain threshold (Figure:4). Recovery of these latencies after IIM suggests that music exerted a genuine antinociceptive effect rather than simply modifying activity levels. The hot plate (HP) and tail flick (TF) tests probe different levels of pain processing, with the TF response being largely spinally mediated and the HP-test engaging more integrated supraspinal processing [12,16]. Therefore, the restoration of both parameters implies that IIM influenced nociception at multiple levels of the neuraxis. Which were further supplemented with the findings from spinal neurotransmitters (Figure:6)and neurotrophic factor (Figure:7) alterations, besides supra-spinal level specific gene-expression studies(Figure:8.1;8.2;9). This is consistent with the hypothesis of rhythmic music engaging descending pain modulatory systems, including the periaqueductal gray and related brainstem circuits, while also affecting cortical and thalamic processing of thermal pain-modulation [26–28].

Thus, the behavioral performances after IIM-exposure reflects a coordinated recovery of sensorimotor function, exploration, and emotional resilience rather than an isolated change in one test parameter. This betterment is important because locomotion, anxiety-like phenotype, and nociceptive thresholds are not independent outputs; they echo reduced ascending pain drive and strengthen endogenous inhibition at different behavioral expressions of a shared pain-stress state.

A major mechanism underlying this behavioral restoration appears to be the normalization of peripheral neurogenic-inflammation. Formalin-pain indicates enhanced primary afferent sensitization and inflammation in paw tissue by elevating the key mediators of neurogenic-inflammation, SP and CGRP which facilitate nociceptor excitability, inflammatory signaling, and NK-1R, the receptor for SP, which participates in dissemination of pain-related neuroimmune cascade[18]. Reduction of them (Figure:5), indicates subsequent attenuation of peripheral nociceptive input at its origin. This regulation was crucial for reducing peripheral afferent nociceptive drive, to further lower both sensory and emotional salience of noxious state, thereby contributing to the restoration of movement, exploratory behavior, and thermal thresholds, ultimately shaping both spinal and supra-spinal responses. Suppression of these biomarkers in music-only group suggests IIM’s tendency toward a basal homeostatic influence, the strongest biological impact was observed when IIM regularize the exaggerated peripheral response in pain-condition.

Pain being a systemic stressor, activates the hypothalamic–pituitary–adrenal (HPA) axis and elevates serum cortisol level indicating substantial stress load, and contributes to anxiety-like behaviors. IIM exposure and reduction of serum cortisol(Figure:9)suggest and strengthens the hypothesis that music dampens stress and interrupts pain-stress conditioning. As pain and stress reinforce each other, so this endocrine modulation leads to neuroendocrinal alteration and then, IIM regulates it at mRNA level of impaired hypothalamic BDNF/Ntrk2R signaling(Figure:9). This subacute pain-stress reflects a maladaptive HPA activation, triggering corticotropin-releasing hormone (CRH) and arginine vasopressin (AVP) eventually leading to elevated serum-cortisol, which ultimately results into impaired trophic signaling at hypothalamic level (reduction of BDNF and Ntrk2) [9].IIM also induces the spinal level expression of dopamine, GABA and lowers glutamatergic signaling reducing stress-reactivity, enhancing reward- and emotion-related systems leading to exploration rather than avoidance. Thus, via normalization of this subacute-stress dysregulated cortisol, hypothalamic BDNF/Ntrk2R signaling; IIM improves the emotional and endocrine pain-stress conditional neurobehaviors; linking peripheral, spinal and eventually the hypothalamic gene-level.

Findings concerning neurochemicals, at the spinal level (L4-L6), acts as a crucial junctional point in between the behavioral recovery, and peripheral effects; being a mechanistic bridge for central nociception. Formalin-pain is associated with excitatory-biased spinal milieu, with elevated level of glutamine, serotonin and reduced GABA, dopamine[10]. This leads to spinal excitatory-inhibitory (E-I) imbalance, resulting into central sensitization and increased dorsal horn excitability. Elevation of spinal glutamate level, activates glutamatergic neurons at spinal dorsal horn which amplifies NMDA/AMPA-driven pain-perception[29] and increased serotonin carries pain-signals via acid-sensing ion channels-3 (ASIC3)[30] leads to hyperalgesia in inflammatory-pain models[31,32]. Gamma-aminobutyric acid (GABA) being a major inhibitory neurotransmitter in the central nervous system (CNS), dampens excessive neuronal excitability [33]. Reduced GABAergic activity implicates neuropathic pain and anxiety disorders [34], formalin also lowered spinal GABA and disrupted the inhibitory control, leading to heightened nociceptive sensitivity [35]. This further supports studies showing that music enhances GABAergic signaling, promoting analgesia and anxiolysis [34]. The lack of significant changes in the music-only group compared to control, suggests IIM’s effect on GABA is context-dependent, requiring a pain-induced imbalance for modulation. 14 days daily music listening reestablished this E-I balance (Figure:6), by lowering excitatory neurotransmitters (glutamine, serotonin) and increasing the inhibitory ones (GABA, dopamine). Our study, provides integrated evidence on dopamine (DA). Formalin-induced subacute-pain triggers a global collapse of the dopaminergic system, characterized by depletion of DA neurotransmitter in spinal-level, simultaneously a concurrent downregulation of D1R mRNA in supraspinal hubs; which signifies subacute-pain induces a “two-hit” hypodopaminergic state [36,37] that negatively affects motivated behaviors, sensory thresholds and mesolimbic-reward system [37]. This depletion is a hallmark of the transition to pathological pain states [36]. Restoration of spinal DA levels suggests that music intervention re-activates descending dopaminergic pathways, effectively “closing the gate” on ascending nociceptive signals in the spinal dorsal horn[38]. Concurrently, the sharp decline in cortical D1R mRNA indicates a failure of the D1R signaling essential for top-down inhibition of hyperexcitable pyramidal neurons[37,39]. Reduction of D1Rs is characterized as, a pro-nociceptive, hyperexcitable state which, effectively lowers the mechanical sensitivity threshold and exacerbates pain perception [39,40]. Increase in the thalamic D1R mRNA after music intervention suggests a modulation of the thalamocortical pathway [37], representing sensory filter mechanism where music-induced DA release reduces the relay of nociceptive information to higher cortical centers. The hippocampal D1R reduction correlates with elevated pain-related anxiety, depression-like states common in persistent pain models [41].

Supraspinal, region-specific significant restoration of D1R, reflects activity-dependent plasticity[42], allows re-establishment of inhibitory control by reactivating reward-linked, top-down modulatory circuits [39,37]. This dopaminergic signaling influences locomotor activation, improved decision-making, and better sensory filtering. Thus, genomic and behavioral data align to show music may re-balance the negative salience of pain with positive reward-related signaling. Interestingly, the music-only group has showed selective changes in D1R expression, indicating that rhythmic auditory stimulus can engage dopaminergic pathways, even in the absence of pain, but the corrective measure can be found in presence of a pathological imbalance.

Spinal as well as supraspinal BDNF emerges as a crucial link in this pain-induced maladaptive state. Although BDNF generally promotes neuronal survival and neurogenesis, BDNF-TrkB signaling, is intricately linked to both adaptive and maladaptive plasticity [43], it also possesses a potent capacity to alter pain pathways at every level of the neuroaxis, from peripheral nociceptors to spinal-neurons eventually brain [44]. At spinal level, BDNF protein upregulation serves as the molecular nexus linking E-I recalibration to upstream relief. Initial formalin-induced BDNF modestly enhances from afferent/microglial sources—typically pro-sensitizing via TrkB-NMDAR crosstalk in acute phases. TrkB engagement traffics KCC2, restoring low intracellular Cl for hyperpolarizing GABA currents and countering depolarization-induced disinhibition. Thus BDNF-KCC2 axis directly bridges neurotransmitter shifts, as enhanced Cl extrusion potentiates phasic inhibition against glutamatergic barrages, while dopaminergic synergy stabilizes interneuron excitability[45]. Thus, the spinal BDNF alteration suggests, IIM did not merely suppress nociceptive activity but promoted adaptive spinal plasticity bridging peripheral and behavioral observations by showing how reduced nociceptive drive was translated into a restored internal gating environment.

BDNF’s supraspinal gene-expression unveiled region-selective BDNF/Trk symphony, delineating functional specialization, and have provided our music-induced pain-management study a genomic dimension with accompanied behavioral and spinal improvements. Formalin-pain altered expression of BDNF, Ntrk1R, and Ntrk2R, across cortex, thalamus, hippocampus, and hypothalamus. The hippocampal and hypothalamic changes crucially map onto the stress– emotion axis that underlies the affective dimension of pain. Hippocampal BDNF, Ntrk1R (TrkA), and Ntrk2R (TrkB) recoveries signals neurogenesis and synaptic remodeling, forging limbic-auditory circuits responsiveness to IIM. This surge recalibrates threat encoding, pain from amygdala-driven fear while amplifying periaqueductal gray (PAG) command neurons[26]. Thalamus gates sensory relay and attentional filtering, and the cortex integrates cognitive and affective appraisal. Thus, the differential expression of Cortical and thalamic BDNF, significantly underlie improved sensory filtering and top-down regulation of nociception. Cortical patterns reflect sensory-emotional integration in anterior cingulate/prefrontal relays, where BDNF/Trk restoration enables cognitive reframing of aversive salience. Thalamic moderation—BDNF/TrkA upregulation with TrkB stability—fine-tunes ventrobasal relay nuclei, filtering somatosensory amplification without overcompensation. Collectively, hippocampal vigor drives PAG-RVM descending amplification, cortical modulation overlays affective overlay, and thalamic gating ensures discriminative precision, converging on brainstem orchestration. These integrates neurobehavioral phenotypes, where antinociceptive thresholds and anxiolytic exploration precisely mirror molecular hierarchies. Thermal reflex recovery tracks spinal E-I gating and PAG-RVM volleys, while locomotor and/or exploratory normalization aligns with hippocampal BDNF-driven affective decoupling, evidenced by open-arm persistence. Music-only enhancements affirm BDNF/Trk’s role in preempting sensitization, establishing rhythmic music as a proactive neuromodulator.

The novelty of this study lies in the integration of our findings supporting, a coherent mechanistic sequence where IIM exposure reduces stress, suppresses peripheral neurogenic-inflammation, restores spinal neurotransmitter and BDNF balance, recalibrates supraspinal gene expression, and ultimately improves pain behavior. Rather than acting as a simple distractor, Indian instrumental music appears to function as a biologically active neuromodulator that recalibrates the stress–nociception axis at multiple levels. In doing so, it improves locomotion, reduces anxiety-like behavior, and normalizes nociceptive thresholds in subacute inflammatory pain. These findings strengthen the promise of rhythmic music as a non-invasive, non-pharmacological pain-management strategy.

## Data Availability Statement

The supporting datasets underlying the results of this study are available from the corresponding author upon reasonable request.

## Declarations

### Funding and Acknowledgment

SD acknowledges the support of funding by DSTBT, Govt. of West Bengal, India, under grant 1804 (Sanc.)/ST/P/S&T/9G-9/2019 and support from DRDO-LSRB, Govt. of India, under grant LSRB/01/15001/M/LSRB-378/SH&DD/2020. SD also acknowledges funding support from ICMR, Govt. of India, under grant 55/02/2022-PHY/BMS. The authors also acknowledge Noah.ai, as it was used for preparation of graphical abstract.

### Contribution of Authors

KM, TB, and SD conceptualized and designed the research; KM, SG, TB performed major experiments using animals; SP, RD, HM performed independent repetition experiments; KM, TB, SG, SP, RDS and SD analyzed results, executed major intellectual input, drafting and editing of the MS.

