## Supplementary for "Melodic Modulation of Pain and Cognition: Neurobehavioral Effects of Indian Classical Music in Mice"

**SUPPLEMEMTARY	DATA FILE**

1.

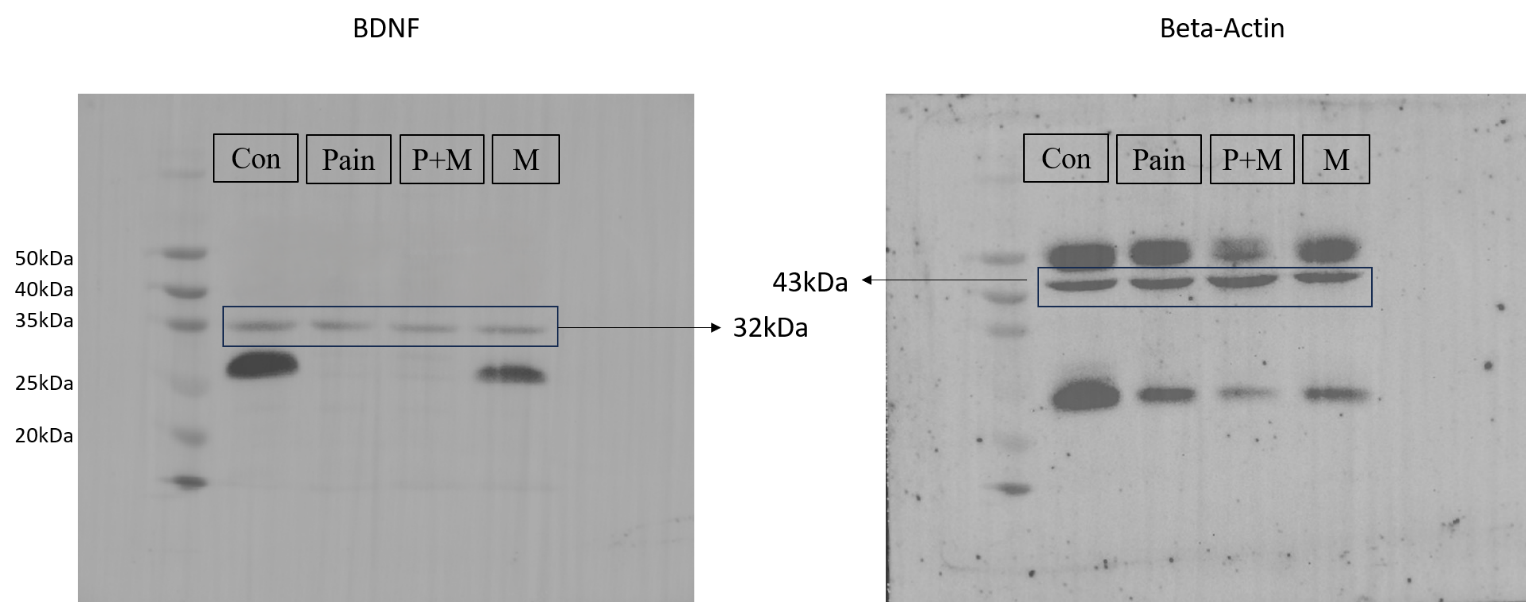

FIGURE 1: Whole Blot images of western Blot of BDNF (A) and stripped Beta-actin in the same blot (B)

2.

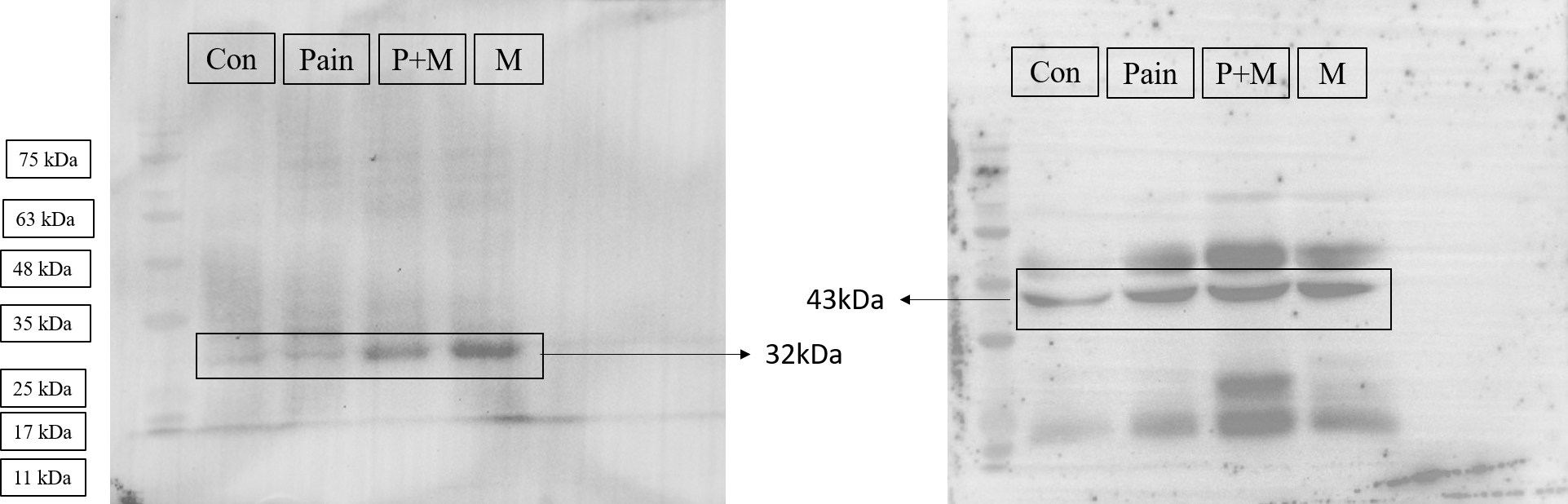

FIGURE 2: Whole Blot images of replicated western Blot of BDNF (A) and stripped Beta-actin in the same blot (B)

3.

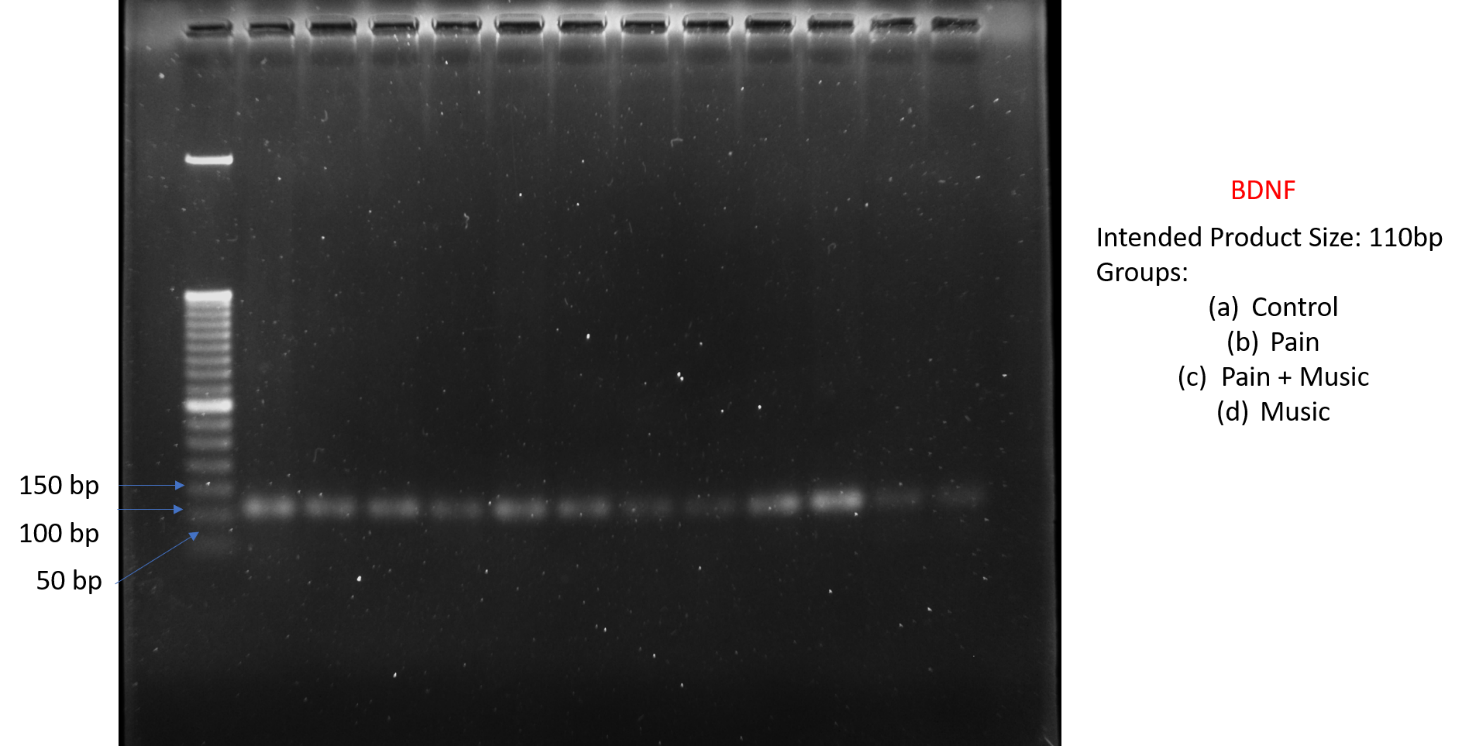

Figure 3: Whole agarose gel image of BDNF gene PCR product at cortex, thalamus, hippocampus across the groups.

4.

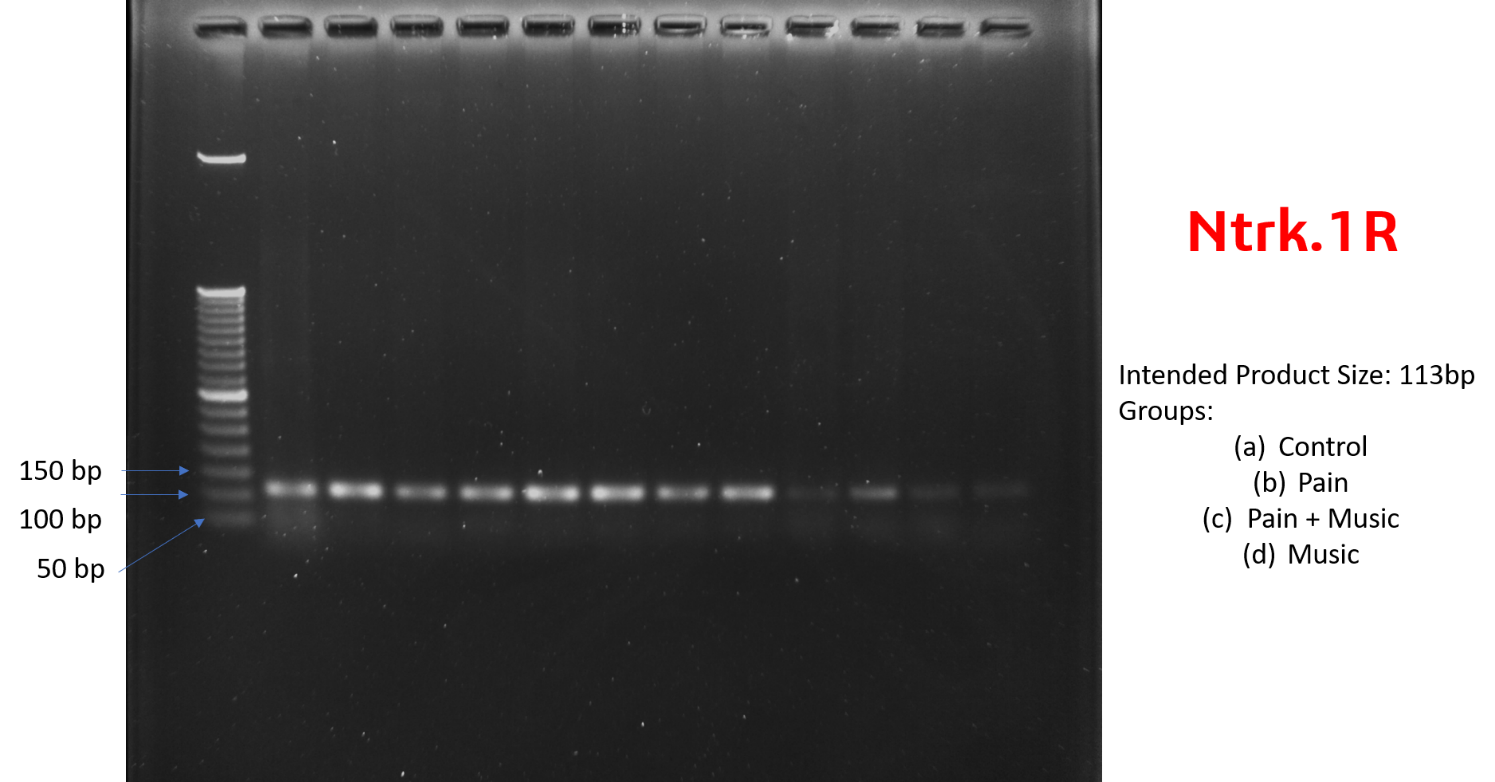

Figure 4: Whole agarose gel image of Ntrk.1R gene PCR product at cortex, thalamus, hippocampus across the groups.

5.

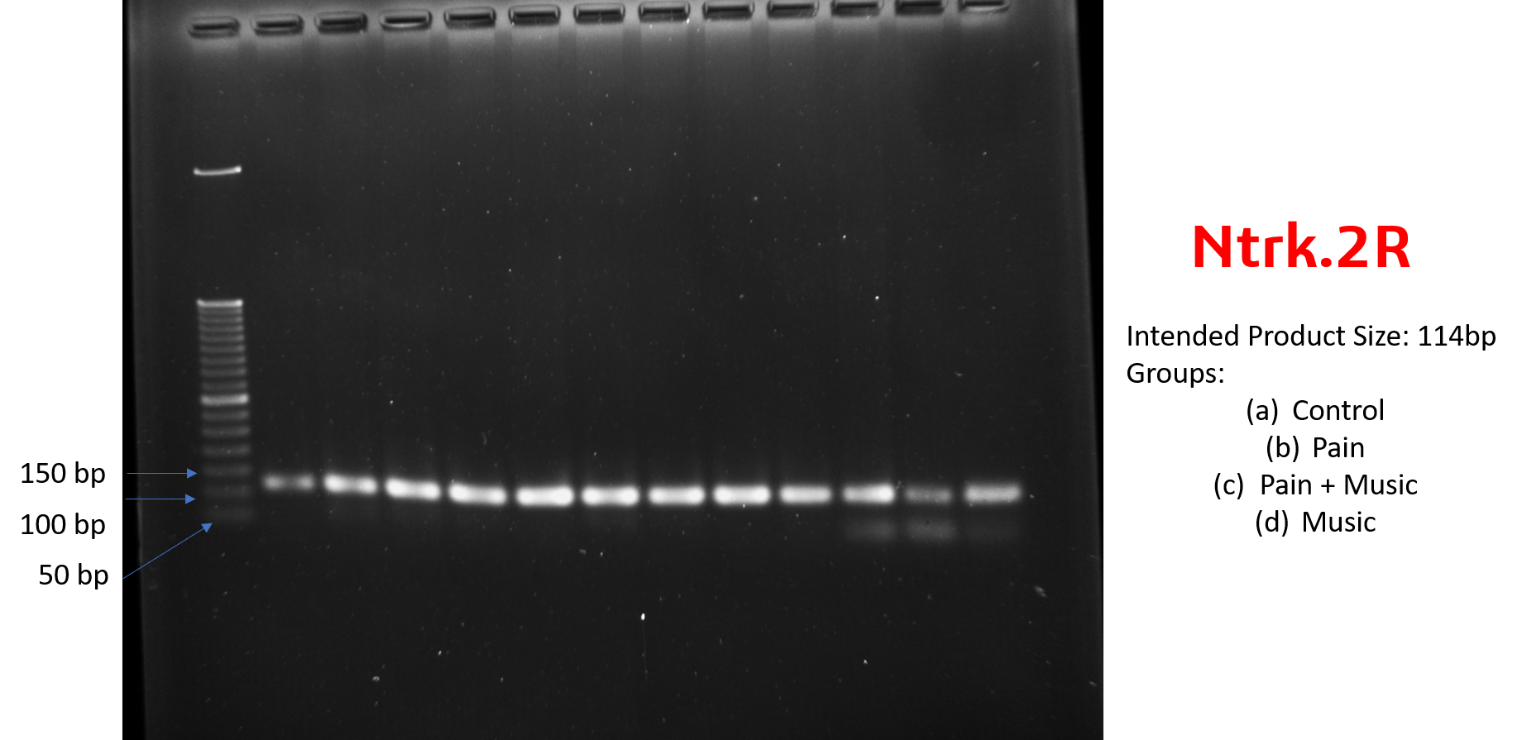

Figure 5: Whole agarose gel image of Ntrk.2R gene PCR product at cortex, thalamus, hippocampus across the groups.

6.
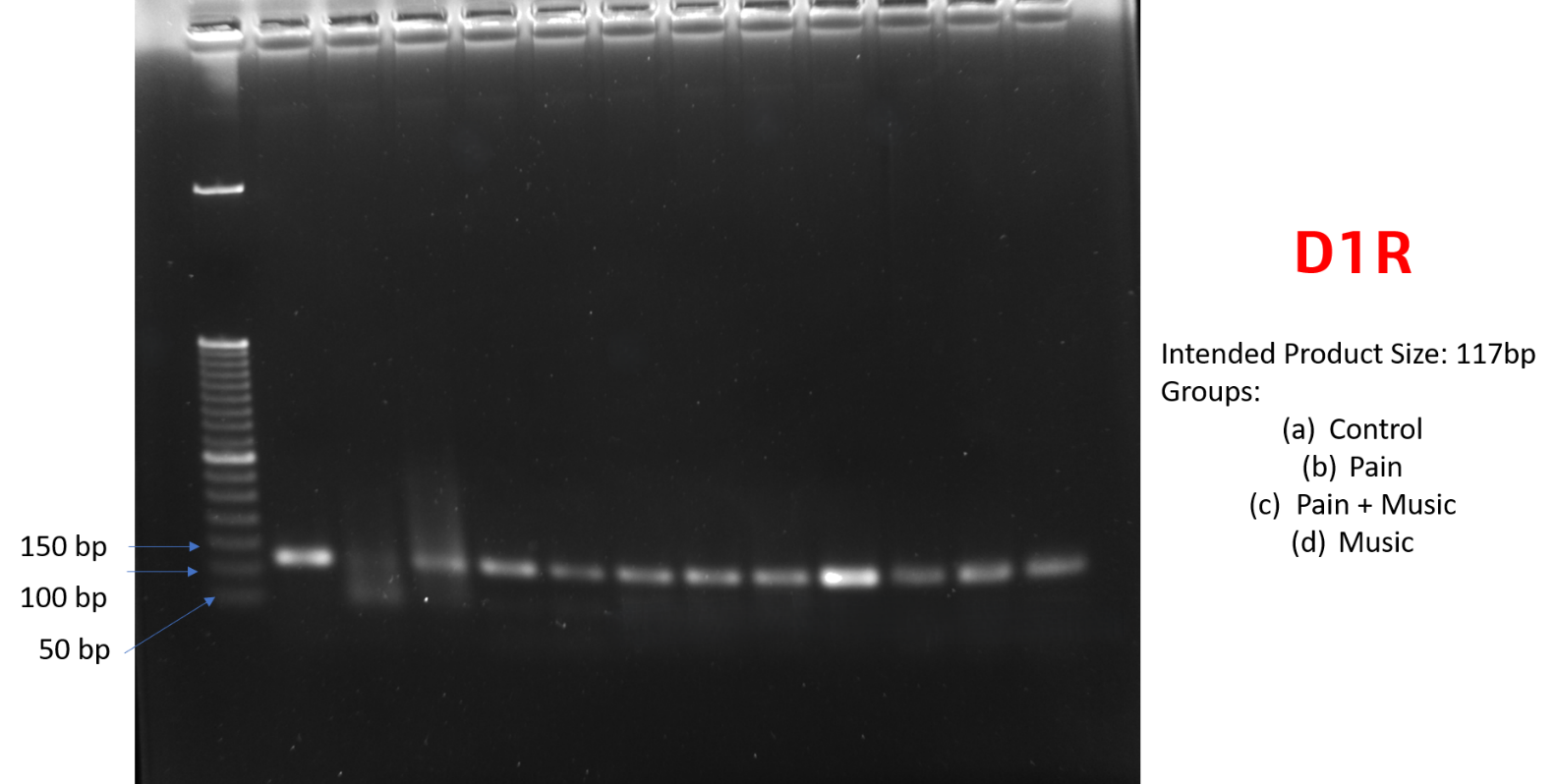

Figure 6: Whole agarose gel image of D1R gene product at cortex, thalamus, hippocampus across the groups.

7.

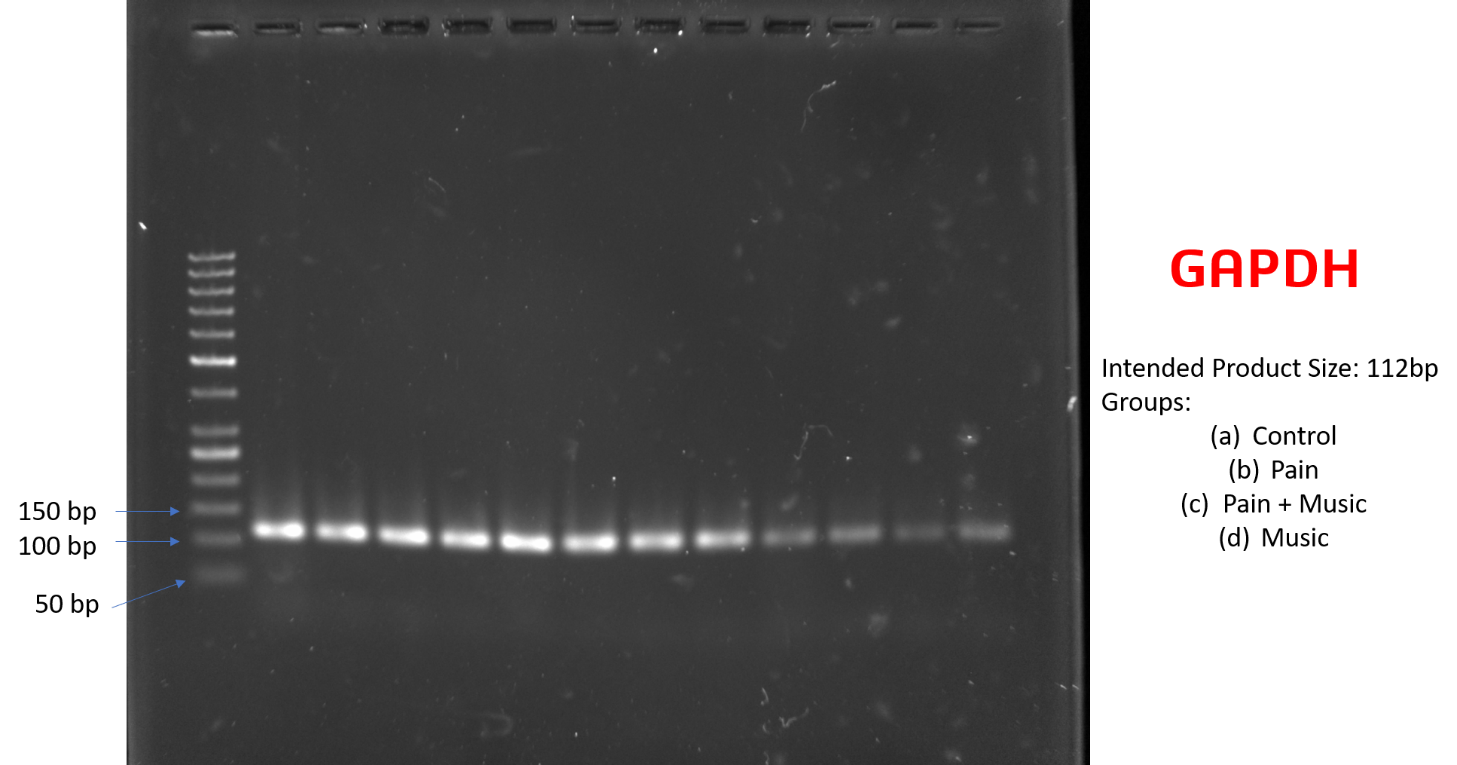

Figure 7: Whole agarose gel image of GAPDH gene PCR product at cortex, thalamus, and hippocampus across the groups.

8.

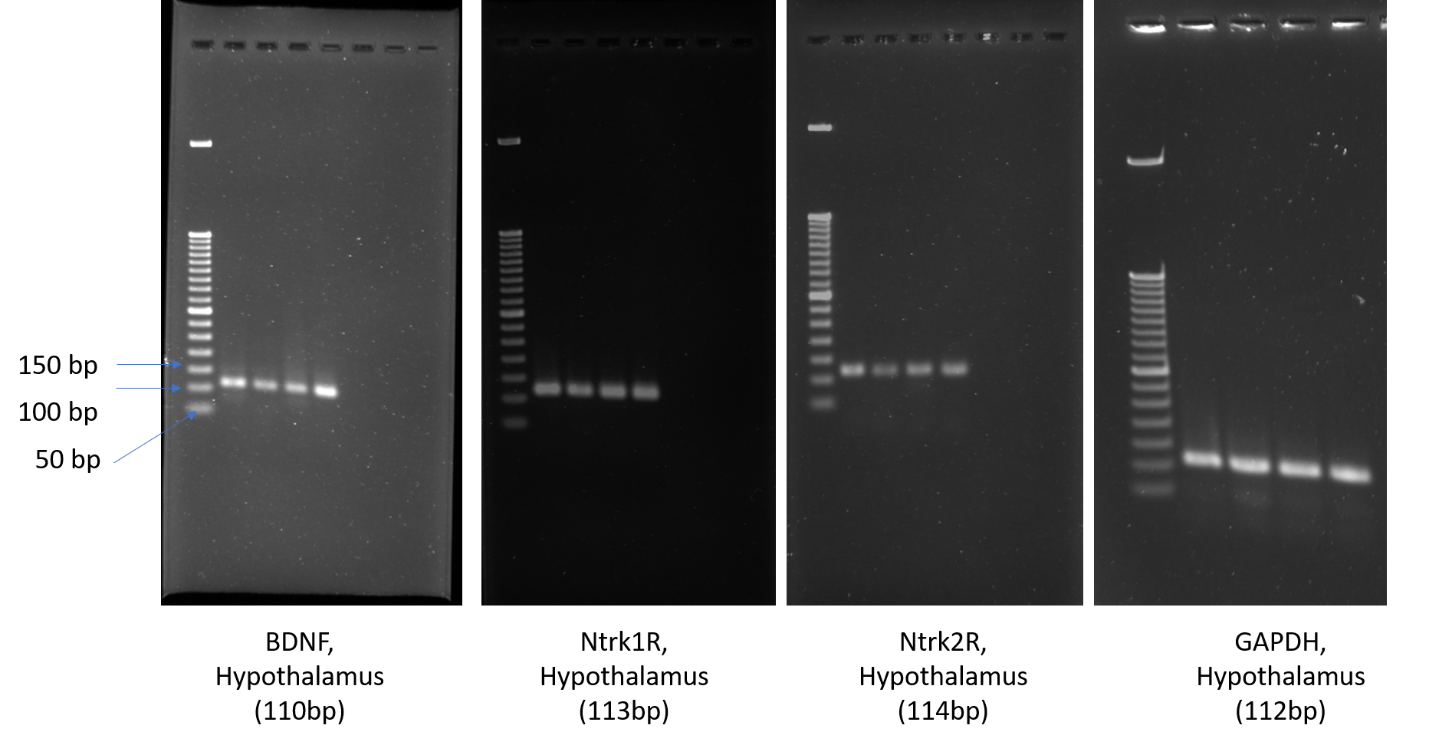

Figure 8: Whole agarose gel image of BDNF, NTRK.1R, NTRK.2R, GAPDH gene PCR product of a groups across in Hypothalamus.

9. **PRIMER TABLE:**

| **GENE NAME** | **FP Sequence** | **RP Sequence** | **Product Size** |
| --- | --- | --- | --- |
| **BDNF** | CAGGGGCATAGACAAAAGGC | TCCTTATGAATCGCCAGCCA | **110 bp** |
| **Ntrk1R** | TGGGGCTGATTCTGGTCAAT | GAAGGAGACGCTGACTTGGA | **113 bp** |
| **Ntrk2R** | CCGGCTTAAAGTTTGTGGCT | CAAGGTGGCGGAAATGTCTC | **114 bp** |
| **D1R** | AGGCTCCATCTCCAAGGACT | GACAGCTTCTCCAGTGGCTT | **117 bp** |
| **GAPDH** | AGTATGACTCCACTCACGGC | ATGTTAGTGGGGTCTCGCTC | **112 bp** |

11.

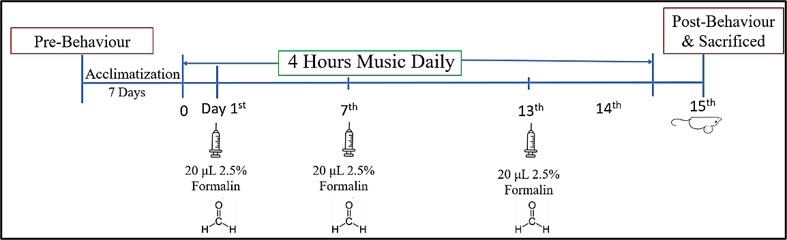

**Figure 1: Study design for 14 days Music exposure**

10. Due to strict regulations of Institutional Animal Ethics Committee (IAEC), University of Calcutta, Kolkata, India; we were not approved to do this study taking more than 3 animals per group;

Keeping this limitation in mind to satisfy the ARRIVE guidelines retrospectively and prove our study mathematically powered to detect biological phenomena, we have calculated the Cohen’s d effect size for all outcomes, comparing the pain group versus the control mice. As Cohen’s d measures the standardized magnitude of an effect independent of sample size. In statistical theory, any effect size exceeding d=0.800 is categorized as large. Our entire experimental effect size demonstrates that this n=3 study, possesses sufficient mathematical power to reliably capture these robust biological transitions. Following is the tabular representation for all the outcomes

**Tabular Characterization of Choen’s d Effect Size (Pain vs. Control):**

| **Biological Domain** | **Specific Outcome Measure Parameter** | **Mean Difference (MD)** | **Pooled Standard Deviation (SD)** | **Calculated Cohen's d (Magnitude)** | **Standardized Effect Size Class** |
| --- | --- | --- | --- | --- | --- |
| **BEHAVIORAL** |  |  |  |  |  |
| · Nociceptive Reflex | Paw Withdrawal Latency (s) | -8.533 | 0.864 | **9.875** | Massive Effect |
|  | Tail Flick Latency (s) | -8.533 | 2.290 | **3.726** | Massive Effect |
| · OFT Anxiety/Locomotor | Center Zone Entries Ratio | -0.717 | 0.175 | **4.087** | Massive Effect |
|  | Center Zone Time Spent Ratio | -0.859 | 0.192 | **4.469** | Massive Effect |
|  | Distance Travelled (DT) Ratio | -0.762 | 0.057 | **13.458** | Massive Effect |
|  | Mean Speed (MS) Ratio | -0.762 | 0.057 | **13.458** | Massive Effect |
|  | Time Immobile Ratio | 3.767 | 1.327 | **2.840** | Massive Effect |
|  | Freezing Time Ratio | 2.741 | 1.170 | **2.343** | Massive Effect |
| · EPM Anxiety-like | Open Arm Time Spent Ratio | -0.731 | 0.074 | **9.840** | Massive Effect |
|  | Open Arm Entries Ratio | -0.990 | 0.190 | **5.214** | Massive Effect |
| **PERIPHERAL TISSUE** |  |  |  |  |  |
| · Paw Lysate | Substance P Concentration | 12.140 | 1.415 | **8.580** | Massive Effect |
|  | CGRP Concentration | 23.810 | 7.436 | **3.202** | Massive Effect |
|  | NK-1R Concentration | 2.373 | 0.187 | **12.724** | Massive Effect |
| **SYSTEMIC** |  |  |  |  |  |
| · Endocrine Axis | Serum Cortisol Level | 7.557 | 1.631 | **4.633** | Massive Effect |
| **SPINAL CORD** |  |  |  |  |  |
| · Neurochemistry | Glutamate Concentration | 1.338 | 0.149 | **8.992** | Massive Effect |
|  | Serotonin Concentration | 0.830 | 0.124 | **6.721** | Massive Effect |
|  | Dopamine Concentration | -1.150 | 0.140 | **8.197** | Massive Effect |
|  | GABA Concentration | -2.377 | 0.210 | **11.319** | Massive Effect |
| · Protein Expression | BDNF Protein (L4–L6 Lysate) | -0.120 | 0.012 | **10.265** | Massive Effect |
| **SUPRASPINAL** |  |  |  |  |  |
| · Cortex mRNA | BDNF Expression | 0.286 | 0.096 | **2.986** | Massive Effect |
|  | Ntrk1R Expression | 0.742 | 0.225 | **3.302** | Massive Effect |
|  | Ntrk2R Expression | 1.849 | 0.090 | **20.590** | Massive Effect |
|  | D1R Expression | -0.455 | 0.032 | **14.128** | Massive Effect |
| · Thalamus mRNA | BDNF Expression | 0.431 | 0.226 | **1.907** | Massive Effect |
|  | Ntrk1R Expression | 1.822 | 0.291 | **6.253** | Massive Effect |
|  | Ntrk2R Expression | 2.510 | 0.973 | **2.581** | Massive Effect |
|  | D1R Expression | -0.093 | 0.008 | **11.675** | Massive Effect |
| · Hippocampus mRNA | BDNF Expression | -1.132 | 0.222 | **5.101** | Massive Effect |
|  | Ntrk1R Expression | 0.840 | 0.265 | **3.170** | Massive Effect |
|  | Ntrk2R Expression | 2.662 | 1.061 | **2.505** | Massive Effect |
|  | D1R Expression | -0.406 | 0.016 | **25.576** | Massive Effect |
| · Hypothalamus mRNA | BDNF Expression | -0.542 | 0.031 | **17.228** | Massive Effect |
|  | Ntrk1R Expression | -0.096 | 0.070 | **1.370** | Large Effect |
|  | Ntrk2R Expression | -0.669 | 0.157 | **4.277** | Massive Effect |

11.

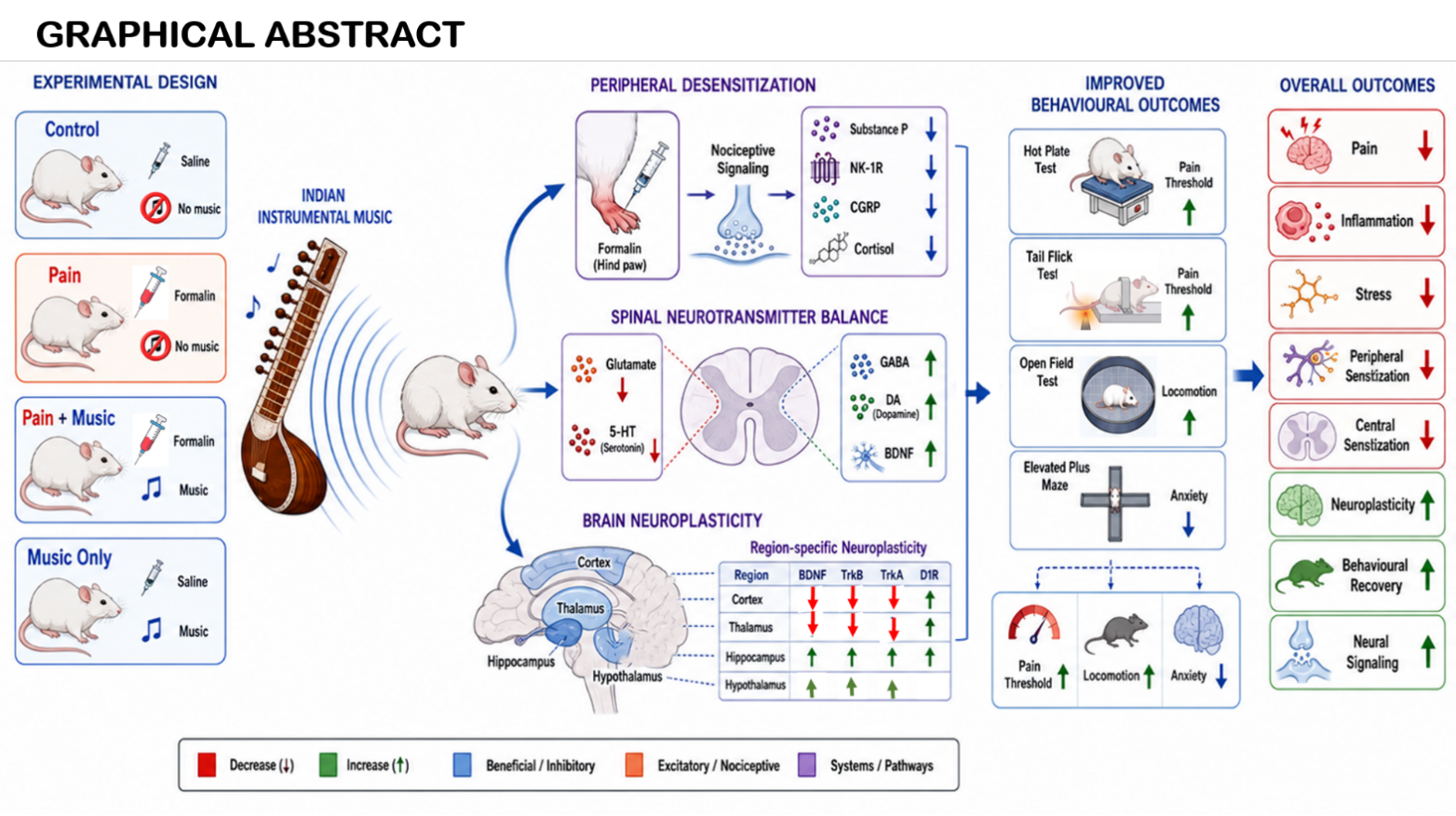
